# Sustained Delivery of a SARS-CoV-2 Subunit Vaccine in An Adjuvanted Hydrogel Depot Enhances Vaccine Responses in Nonhuman Primates

**DOI:** 10.64898/2026.07.17.736840

**Authors:** Ben S. Ou, Mengyun Hu, Jerry Yan, Jordan M. Santagata, Olivia M. Saouaf, Haleigh B. Eppler, Victor Lujan, Ye Eun Song, Alba Grifoni, John H. Klich, Alessandro Sette, Ashley Utz, Mehul S. Suthar, Noah Eckman, Yupeng Feng, Julie Baillet, Kenneth A. Rogers, Lisa M. Shirreff, George J. Aoyagi, Adian S. Valdez, Rashmi Ravichandran, Neil P. King, Jane Fontenot, Francois Villinger, Bali Pulendran, Eric A. Appel

**Author notes:** These authors contributed equally to this work (Ben S. Ou, Mengyun Hu). Corresponding Authors &.

## Abstract

While natural infections expose the immune system for days to weeks of inflammation and antigen presentation, immunizations with conventional bolus vaccines often lead to rapid clearance of antigens and adjuvants. Prolonged exposure to vaccines using controlled delivery devices or repeated dosing regimens has been shown to enhance germinal center reactions, leading to improved humoral responses, including increased magnitude of antibody titers and enhanced neutralizing activity. Herein, we report the use of injectable polymer-nanoparticle (PNP) hydrogels as a vaccine depot technology for sustained delivery of the clinically relevant SARS-CoV-2 Hexapro subunit antigen and a toll-like receptor agonist adjuvant. In mice, we demonstrated that PNP hydrogel vaccines enhanced germinal center responses and antibody responses relative to bolus vaccination. In nonhuman primates, hydrogel vaccines induced enhanced and durable antibody responses against wildtype and variants of concern such as Omicron BA.5 compared to bolus vaccination. We report the first use of a biomaterials-based approach for sustained delivery of vaccines in nonhuman primates, further advancing toward clinical translation.

## Introduction

Vaccines have been one of the greatest medical interventions of the past century, and the emergence of global pandemics such as human immunodeficiency virus (HIV), influenza, and the most recent severe acute respiratory syndrome coronavirus 2 (SARS-CoV-2) have further catalyzed their development. Yet vaccines that provide durable and broad protection against these rapidly mutating viral pathogens have yet to be developed.^1^ Indeed, while the original SARS-CoV-2 mRNA/LNP vaccines were highly effective against the wildtype variant, they were shown to lose 20-30% of their efficacy within 6 months and were only ∼8.8% effective against the highly resistant Omicron variant.^2–7^ Similarly, no cross-reactive or broadly neutralizing antibodies (bnAbs) against HIV have to date been detected in the sera from humans or nonhuman primates (NHPs) following vaccinations with the most promising HIV vaccine candidates.^8^

Since high-affinity and broadly neutralizing antibodies are believed to arise from affinity maturation and somatic hypermutation (SHM), highly active and prolonged germinal center (GC) responses may be pivotal in enhancing protection against these difficult-to-target antigens.^9–23^ Recently, the control over vaccine exposure kinetics through sustained release of antigen and adjuvant during vaccination has been shown to improve humoral responses.^22, 23^ The sustained exposure of HIV vaccines in NHPs over 7-28 days using osmotic pumps or repeated injections has been shown to prolonged germinal center reactions and enhance immune responses compared with traditional bolus immunizations.^12, 13, 15^ However, surgically implanted devices or the need for multiple injections over a week or two for vaccinations remain challenging to translate into clinical settings, particularly in low-resource settings. While aluminum salts (Alum) and other adjuvants have been believed to act as depots for the sustained delivery of antigens, many studies have reported rapid antigen elution from Alum and overall inefficiency in improving antibody responses *in vivo*.^24–29^ Therefore, a more effective biomaterial-based method for sustained vaccine delivery is necessary for clinical application.

In this context, we have previously reported the use of an injectable polymer nanoparticle (PNP) hydrogel depot technology that enables sustained co-delivery of diverse antigens and adjuvants for up to 4 weeks, mimicking the exposure kinetics of natural infections.^30–41^ We showed that these PNP hydrogels enable robust, durable, and broad immune responses to vaccines against influenza and SARS-CoV-2 in murine models.^30–41^ Here, we leveraged the PNP hydrogel platform for encapsulation and sustained delivery of SARS-CoV-2 vaccines comprising the clinically relevant Hexapro antigen and the TLR7/8 agonist adjuvant 3M-052^42–45^ in both mouse and nonhuman primate models. We first characterized the mechanical properties of different PNP hydrogels for sustained co-delivery of antigen and adjuvant. We then showed that mice vaccinated with PNP hydrogel Hexapro vaccines exhibited enhanced germinal center responses, higher binding and neutralizing titers, and enhanced memory responses compared with an Alum bolus control. In a nonhuman primate model, we further demonstrated that PNP hydrogel vaccines increased the magnitude and durability of antibody binding titers, enhanced germinal center responses, enriched CD4+ T cell and memory B cell responses, and broadened responses against variants of concern. Together, we established that PNP hydrogel vaccines are promising clinical candidates as biomaterials for slow-release vaccination to enhance humoral responses (Figure1a).

## Results

### Formulation of PNP Hydrogels for Sustained Delivery of Hexapro Vaccines

PNP hydrogels are formed rapidly by mixing aqueous solutions of dodecyl-modified hydroxypropylmethylcellulose (HPMC-C_12_) with biodegradable poly(ethylene glycol)-poly(lactic acid) nanoparticles (NPs) (Figure 1b). Entropy-driven polymer-nanoparticle interactions between the HPMC-C_12_ chains and the NPs produces physically crosslinked hydrogels that are shear-thinning and self-healing.^46, 47^ These properties enable injection through standard-gauge needles and promote rapid formation of a robust depot after injection.^32, 48, 49^ This dynamically crosslinked depot enables sustained co-release of diverse biologic cargo over the course of 2-4 weeks, and enables formation of a transient, local inflammatory niche for immune cell infiltration that enhances antigen processing (Figure 1a, 1b).^33, 35, 36, 41^

**Figure 1.**
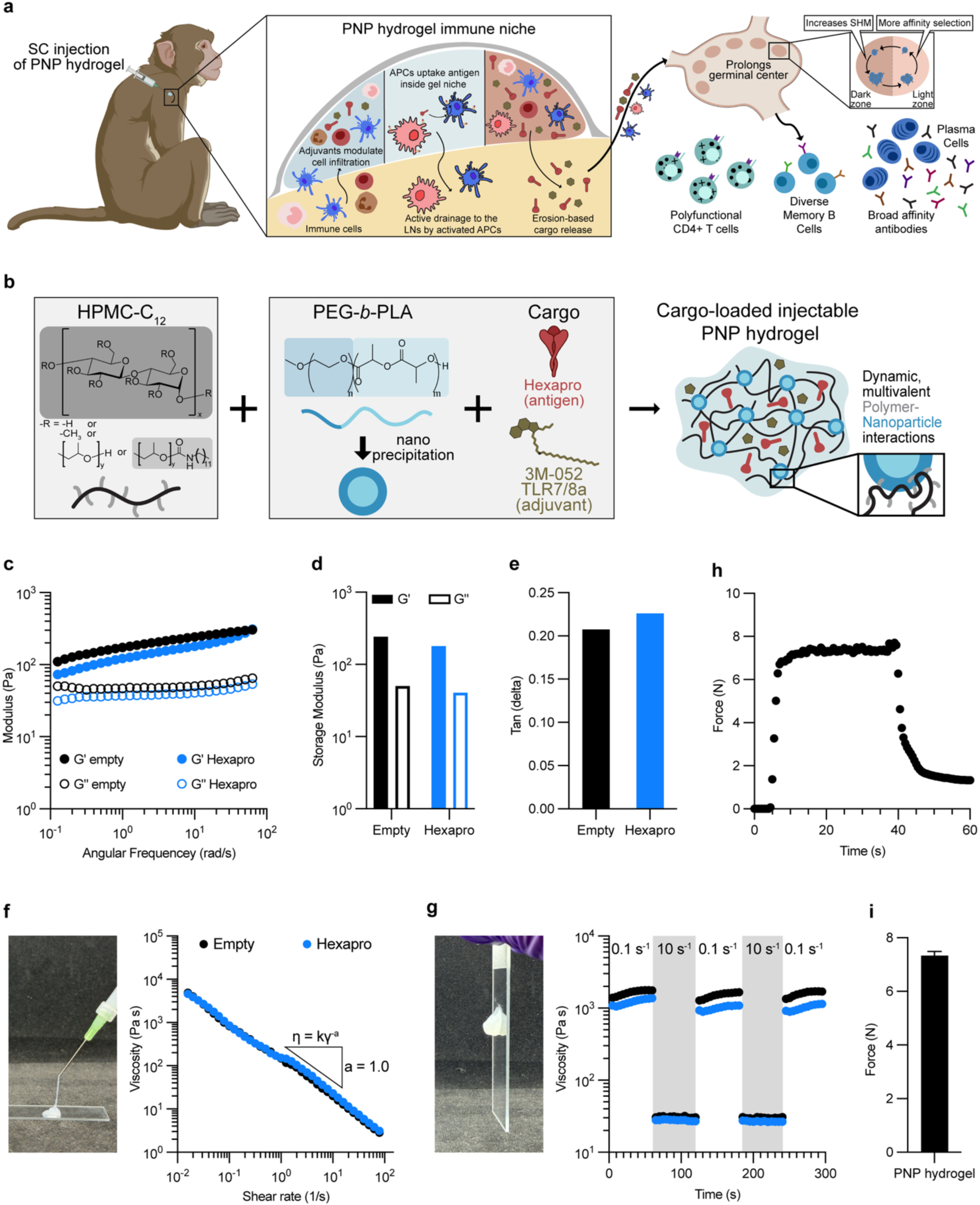
Formulation and Characterization of PNP Hydrogels for Sustained Delivery of Hexapro Vaccines. **(b)** Schematic of mixing HPMC-C_12_ Polymer with PEG-*b*-PLA nanoparticles and vaccine cargoes (SARS-CoV-2 antigen Hexapro and 3M-052 adjuvant) to form vaccine-loaded Polymer-Nanoparticle (PNP) hydrogel vaccines. **(c)** Frequency-dependent oscillatory shear rheology of empty and Hexapro-containing PNP hydrogels. **(d)** Storage (G’) and Loss (G”) moduli and (**e**) Tan (δ) of PNP hydrogel formulations at an angular frequency of ω = 10 rad/s. **(f)** Representative injection image through a 21-gauge needle and shear-dependent viscosities of PNP hydrogels with a large power-law shear-thinning index “a”. **(g)** Representative solid-like depot image and step-shear measurements of PNP hydrogels over three cycles of alternating high shear (10 s^-1^) and low shear (0.1 s^-1^) rates. **(h)** Injection force curves and **(i)** corresponding plateau injection forces (*n* = 3) for injection of PNP hydrogels at 1 mL/min through 21-gauge, 1-inch needles. Results are shown as mean ± SD.

In this study, we sought to use PNP hydrogel platform for the sustained co-delivery of Hexapro antigen and TLR7/8a 3M-052 adjuvant. We showed that the viscoelastic properties of PNP hydrogels with the presence of Hexapro and 3M-052 were unaffected compared to empty PNP hydrogels, exhibiting similar stiffness and Tan(δ) values (Figures 1c-1e). Similarly, shear-thinning property remained unchanged, as evidenced by the large decrease in viscosity with increasing shear rate (with representative injection image, Figure 1f). PNP hydrogels exhibited a shear-thinning index, a, from the relationship η = kγ^-a^ of around 1, indicating a high degree of shear thinning that is necessary for injectability.^50–52^ Furthermore, using our flow rheology data, we modeled the extrudability of the PNP hydrogels with and without cargo and predicted that both formulations would be injectable under clinically relevant pressures and forces (Figure S1).^48, 53^ Self-healing and injectability properties persisted with the addition of cargo since we observed rapid recovery in step shear measurements across multiple cycles when applying stepwise change between low and high share rates (0.1 s^-1^ and 10 s^-1^, respectively; with representative solid-like depot image, Figure 1g). Overall, the inclusion of vaccine components does not compromise the rheological properties of the PNP hydrogels.

We further evaluated the injectability of these PNP hydrogel vaccine formulations by measuring the force required to inject them through a 21-gauge, 1-inch needle to mimic clinical application (Figure 1h). For therapeutic applications, an injection force of less than 30N is required for subcutaneous or intramuscular injections by hand.^54, 55^ At a flow rate of 1 mL/min, the injection force was 7.3 N, well below the clinically relevant maximum threshold (Figure 1i). These studies confirmed that the PNP hydrogels can be used clinically for the sustained delivery of vaccines.

### PNP Hydrogel Hexapro Vaccines Enhanced Germinal Center Responses and Durable Antibody Responses in Mice

We immunized C57BL/6 mice (n = 6) subcutaneously with Hexapro vaccines (0.5 μg) adjuvanted with 3M-052 adjuvant (1 μg), formulated either with Alum (100 μg) or PNP hydrogels. The bolus vaccine with Alum as a depot was used to mimic clinical adjuvant formulations and previous studies (denoted Alum bolus).^31, 41, 43–45, 56–59^ Mice were primed at Week 0 and boosted at Week 8 to replicate clinical schedules. Serum, draining inguinal lymph nodes (dLNs), and bone marrow were collected at periodic intervals for analysis (Figure 2a). We first measured antibody binding responses across vaccine groups using enzyme-linked immunosorbent assays (ELISAs) on serum samples to determine endpoint IgG titers. Mice vaccinated with the PNP hydrogel vaccine produced significantly higher endpoint IgG binding titers than those immunized with Alum bolus at all time points over 24 weeks (p = 0.002 for AUC; Figure 2b, 2c). Notably, all but one mouse vaccinated with PNP hydrogel seroconverted by Week 1, and all mice in this group had detectable titers by Week 2. In contrast, two of six mice immunized with Alum bolus did not seroconvert until after the boost. Furthermore, the highest average titer of the Alum bolus group post-boost (Week 12) was still at least 40% lower than the average titer of the PNP hydrogel group before boost, highlighting the enhanced humoral immune responses generated by the PNP hydrogel vaccines.

**Figure 2.**
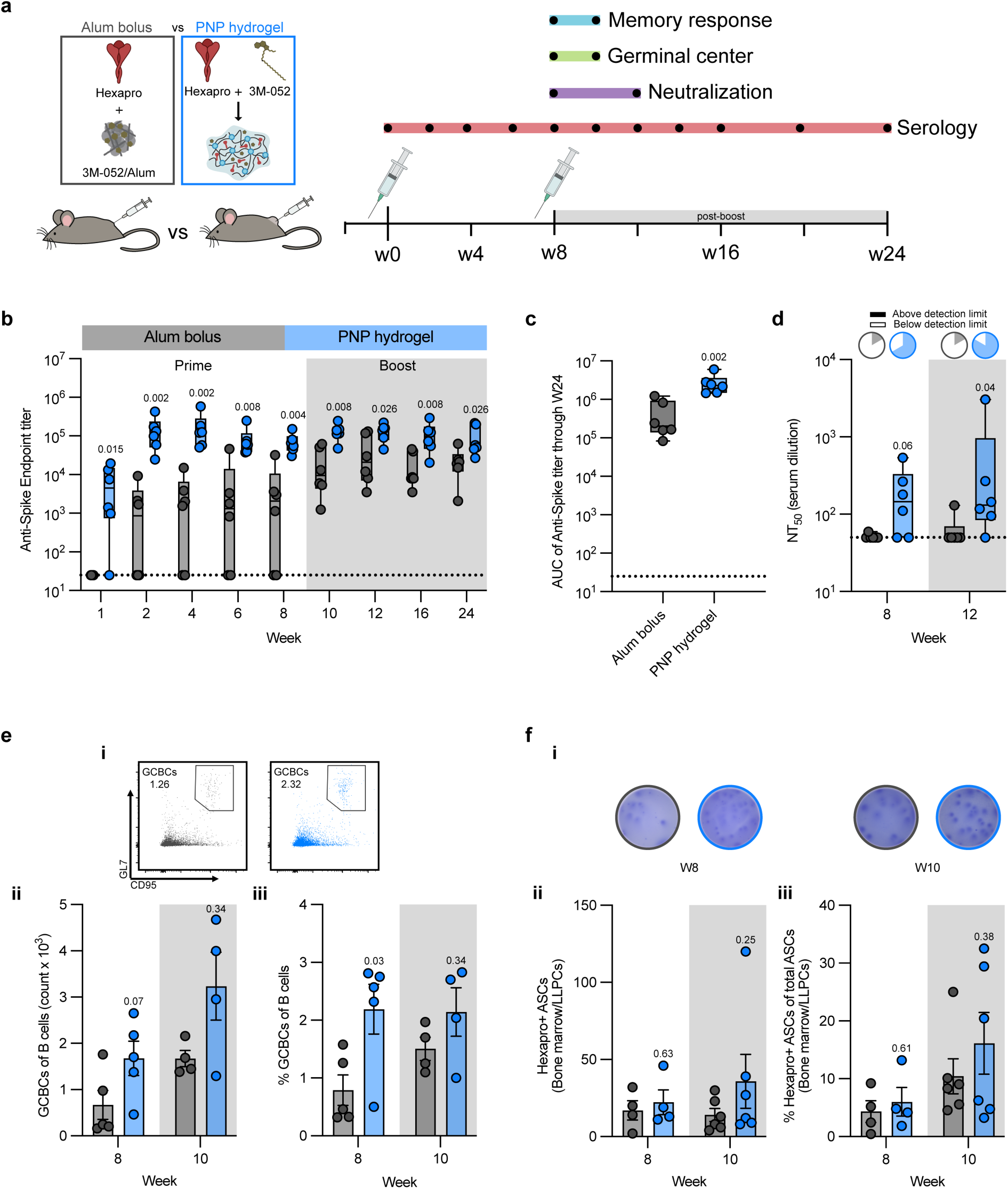
PNP Hydrogel Hexapro Vaccines Induced Robust and Durable Responses in Mice. **(a)** Timeline for immunization and sample collection to assess humoral responses to vaccination. Mice were immunized at Weeks 0 and 8 with either the Alum bolus or the PNP hydrogel Hexapro vaccine, adjuvanted with 3M-052. **(b)** Anti-spike IgG endpoint titers over time. **(c)** Area under the curve (AUC) of anti-spike titers from (b). **(d)** Neutralizing activity of sera from immunized mice against wild-type SARS-CoV-2 pseudoviruses pre- and post-boost (W8, W12) represented as NT_50_ values. Pie charts depicted the ratios of responders to nonresponders at each time point, defined as mice with no detectable neutralizing activity. **(e) (i)** Representative flow plots and quantification of germinal center B cell (GCBC) **(ii)** counts and **(iii)** percentages of all B cells of each vaccine group pre- and post-boost (W8, W10). **(f) (i)** Representative flow plots and **(ii)** counts and **(iii)** percentages of Hexapro-specific antibody-secreting cells (ASCs) from the bone marrow of each vaccine group pre- and post-boost (W8, W10). Data are shown as box-and-whiskers plots showing median, interquartile (box), and range (whiskers) or as bars with mean ± SEM (n = 4-6). The *p* values listed were determined by the unpaired Mann-Whitney test.

We then tested the neutralizing activity of the sera by measuring the serum-mediated inhibition of SARS-CoV-2 spike pseudotyped lentivirus entry into HeLa cells overexpressing ACE2 and TMPRSS2. We tested the neutralizing activity of sera from both groups at Week 8 (pre-boost) and Week 12 (post-boost) across a range of concentrations to determine the half-maximal inhibition of infectivity (NT_50_) (Figure 2d, S2). Only one out of the six mice in the Alum bolus group had detectable neutralizing activity in its serum at either time point. In contrast, all but one mouse in the PNP hydrogel vaccine group generated sera with robust neutralizing activity at Week 12, resulting in higher NT_50 values_ (p = 0.06 at Week 8 and p = 0.04 at Week 12).

To better understand the mechanisms underlying the observed improvements in the potency and durability of the PNP hydrogel vaccine, we used flow cytometry to probe GC responses at Weeks 8 and 10 (Figure 2ei-iii, S3). We observed sustained GC B cell (GCBC) responses following vaccination with PNP hydrogel vaccines, with higher GCBC counts and significantly higher GCBC percentages detected at Week 8 post-prime compared with the Alum bolus group (p = 0.07 and p = 0.03, respectively). Similarly, PNP hydrogel vaccines also induced higher GCBC counts and percentages post-boost (Week 10). We then assessed memory responses by analyzing the population of Hexapro-specific antibody-secreting cells (Hexapro+ ASCs) in the bone marrow using enzyme-linked immunospot (ELISpot) assays, as this population primarily comprises long-lived plasma cells. Mice vaccinated with PNP hydrogel vaccines had higher counts and percentages of Hexapro+ ASCs than those immunized with Alum bolus at both Week 8 and Week 10 (Figure 2fi-iii). At Week 10, more than twice the count of Hexapro+ ASCs was measured following PNP hydrogel vaccination when compared to Alum bolus (p = 0.25). Overall, these results show that vaccination with PNP hydrogels improve humoral responses in mice.

### PNP Hydrogel Hexapro Vaccines Enhanced and Prolonged Antibody Titer Responses in Rhesus Macaques

To build on our murine findings and enhance their translational relevance, we investigated immune responses in rhesus macaques (RMs) using a comparative vaccination study. We immunized RMs (n=6) with Hexapro vaccines (12.5 μg) adjuvanted either with 3M-052/Alum (5 μg 3Μ-052/500 μg Αlum, referred to as Alum bolus) or PNP hydrogel adjuvanted with 3M-052 (5 μg, referred to as PNP hydrogel). The RMs were primed on Week 0 and boosted on Week 8, matching the mouse study and previously reported regimens. Serum, peripheral blood mononuclear cells (PBMCs), and fine lymph node aspirates were collected for analysis (Figure 3a).

**Figure 3.**
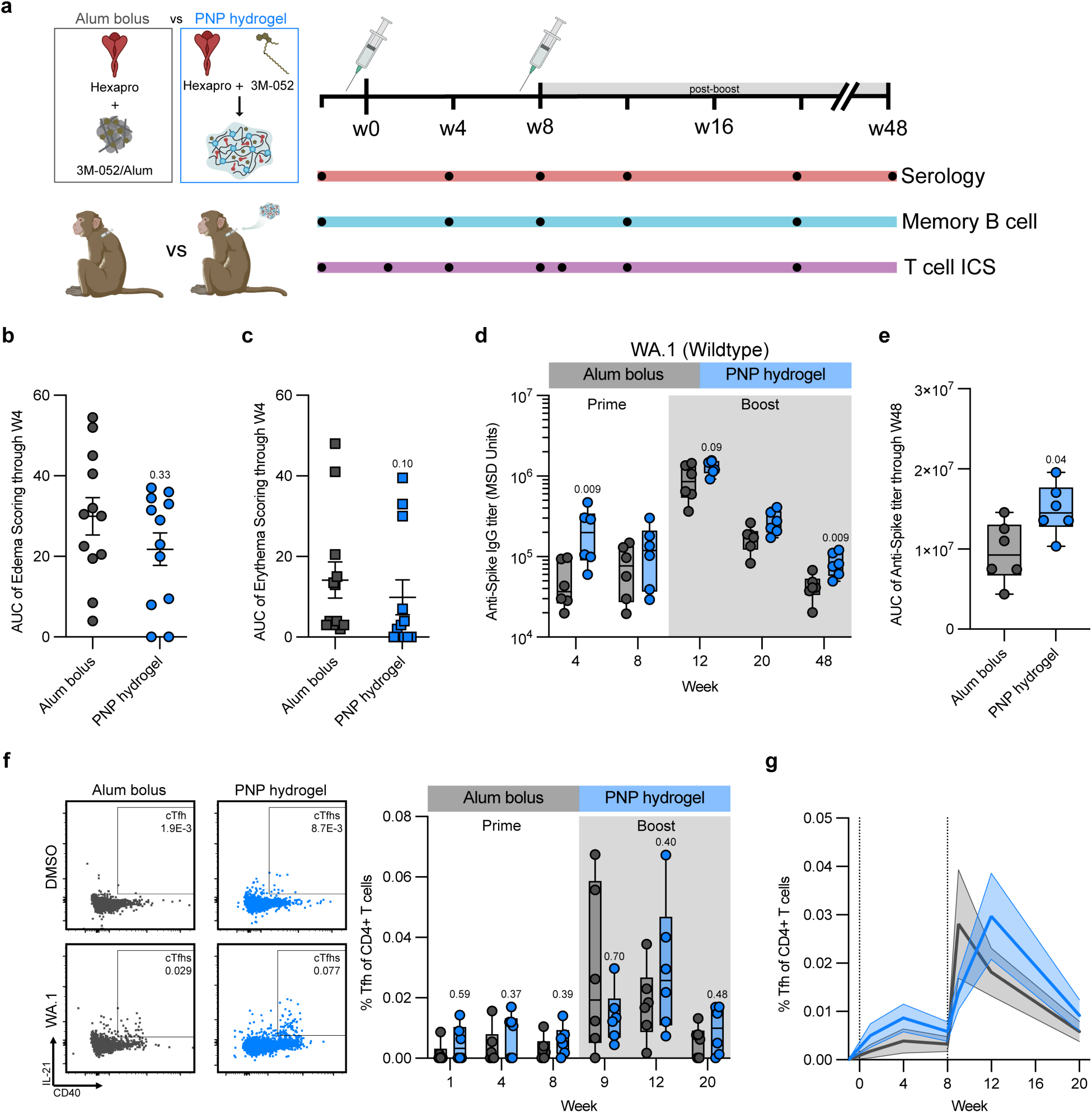
PNP Hydrogel Hexapro Vaccines Enhanced and Prolonged Titer Responses in Rhesus Macaques. **(a)** Timeline for immunization and sample collection to assess humoral responses to vaccination. Nonhuman primate rhesus macaques (RMs) were immunized at Weeks 0 and 8 with either the Alum bolus or the PNP hydrogel Hexapro vaccine, adjuvanted with 3M-052. Area under the curve of **(b)** clinical edema score and **(c)** clinical erythema score of the injection site over 4 weeks. **(d)** Anti-spike IgG titers over time. **(e)** Area under the curve (AUC) of anti-spike titers from (d)**. (f)** Representative flow plots showing expression of IL-21 and CD40 to define circulating T follicular helper (cTfh)-like cells in the blood after ex vivo stimulation with DMSO (no peptide, top) or an overlapping peptide pool spanning the SARS-CoV-2 spike (bottom) and the percentage of spike-specific cTfhs of all CD4+ T cells of each vaccine group over time. **(g)** The spike-specific cTfh CD4+ T cell responses in the blood of each vaccine group, shown as mean and SEM over time. Data are shown as box-and-whiskers plots showing median, interquartile (box), and range (whiskers) or as bars with mean ± SEM (n = 6). The *p* values listed were determined by the unpaired Mann-Whitney test.

As the first reported use of an injectable biomaterial as a depot technology for sustained vaccine delivery in a large animal model, we assessed whether injection-site reactions would occur following PNP hydrogel vaccine injections. Clinical edema and erythema scores were assessed daily for the first two weeks, then weekly until Week 4 (Figure S4a). PNP hydrogel vaccines produced post-injection site reactions similar to those of the Alum bolus control formulation and elicited lower edema and erythema clinical scores on average (based on AUC; p = 0.33 and p = 0.10, respectively; Figure 3b, 3c, S4b, S4c). Thus, the PNP hydrogel vaccine not only did not cause adverse reactions but may also reduce overall reactogenicity by slowly delivering the proinflammatory and immunogenic vaccines comprising 3M-052.

Next, we measured the RMs’ antibody binding titers over time using Meso Scale Diagnostics (MSD) ECL-ELISAs, focusing on the higher-quality IgG antibodies. As early as Week 4, RMs immunized with PNP hydrogel generated robust IgG titers, four-fold higher than those vaccinated with Alum bolus (p = 0.009; Figure 3d). Titers from the RMs vaccinated with PNP hydrogel remained higher at all time points compared with the Alum bolus group, with titers remaining twice as high at Week 48 (p = 0.009), resulting in the PNP hydrogel group with significantly higher AUC titer overall (p = 0.04; Figure 3e). Using this long-term titer data, we assessed the antibody decay half-life from post-boost peak titers using parametric bootstrapping based on an exponential decay model (Figure S5). We found that immunizing with PNP hydrogel significantly increased the half-life by more than 3 weeks compared to immunizing with alum bolus (117 days vs 92 days; p < 0.0001). Thus, PNP hydrogel vaccines enhanced the magnitude and early persistence of titer responses with features consistent with improved durability.

Since enhanced and prolonged antibody titers are likely the result of sustained GC reactions, we therefore sought to determine the differences in GC activities. First, we isolated PBMCs, stimulated them *ex vivo* with a spike-peptide pool, and quantified the circulating-T follicular helper cells (cTfhs; CD40L- and IL-21-double-positive)^60^ populations via flow cytometry since GC activities are mediated by Tfhs and are critical for generating broad and durable vaccine responses. (Figure 3f, S6). While cTfhs from RMs receiving Alum bolus peaked at Week 9, just one week post-boost, cTfhs from RMs receiving the PNP hydrogel peaked at Week 12, four weeks post-boost (Figure 3f, 3g). Measuring the total cTfhs over time by assessing the AUCs, we found that RMs immunized with PNP hydrogels resulted in more than double the total cTfhs compared with immunizing with Alum bolus during prime (AUC of 0.05 vs 0.023, respectively; p = 0.13; Figure S7a) and maintained higher AUC over the whole study period (AUC of 0.29 vs 0.2, respectively; p = 0.39; Figure S7b).

We also examined early GC responses after vaccination using fine-needle aspiration of the draining ipsilateral lymph nodes to assess GCBCs and GC-Tfh populations (Figure S8a). Four weeks post-prime, the levels of both GCBCs and GC-Tfhs from the RMs immunized with Alum bolus vaccines returned to similar levels as the baseline, which were measured 1 week before vaccination (p = 0.13 and p = 0.30, respectively; Figure 8b, 9c). In contrast, GCBCs and GC-Tfhs from those vaccinated with PNP hydrogel remained significantly higher than the baseline levels at Week 4 (p = 0.002 and p = 0.026, respectively; Figure 8b, 8c). These findings suggest the hydrogels’ ability to improve early GC persistence, enhancing the titer responses.

### PNP Hydrogel Hexapro Vaccines Induced Higher Polyfunctional Th1 CD4+ T Cell Responses in Rhesus Macaques

Since antigen-specific B cells require the help of CD4+ T cells, we measured the cellular responses following immunization with either Alum bolus or PNP hydrogel Hexapro vaccines. To determine the durability and potency of CD4+ T cells, specifically Th1 CD4+ T cells, we isolated PBMCs and stimulated them *ex vivo* with an overlapping peptide pool spanning the Spike protein, as previously described.^61–63^ Wildtype spike-specific Th1 CD4+ T cell populations were determined by grouping interleukin-2- (IL-2), interferon-gamma- (IFN-γ), and tumor necrosis factor-alpha-(TNF-α) secreting CD4+ T cells via flow cytometry (Figure 4a, S4). Before boosting, only RMs vaccinated with PNP hydrogel resulted in significant Th1 CD4+ T cell activation, as determined by comparison with pre-immunization levels (p = 0.04, p = 0.19, respectively; Figure 4b).

**Figure 4.**
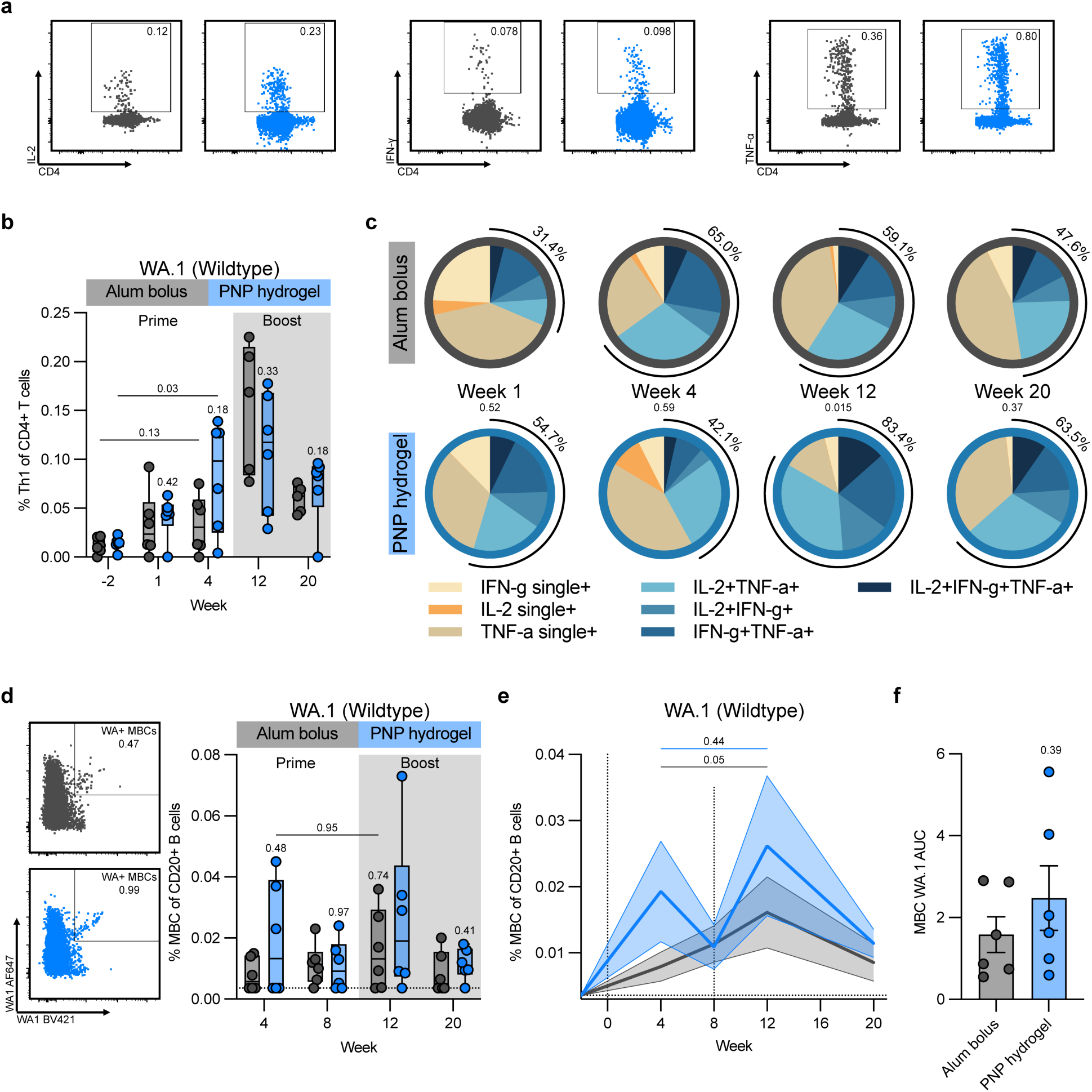
PNP Hydrogel Hexapro Vaccines Induced Higher Polyfunctional CD4 T Cell and Extended Memory B Cell Responses in Nonhuman Primates. **(a)** Representative flow charts of IL-2+, IFN-γ+, and TNF-α+ CD4+ T cells to determine Th1 CD4+ T cell populations measured in blood**. (b)** Percentages of wildtype spike-specific Th1 CD4+ T cells of all CD4+ T cells of each vaccine group over time. **(c)** Pie charts representing the fractions of each monofunctional (single cytokine-specific, beige hues) and polyfunctional (double or triple cytokine-specific, blue hues and outlined in black) Th1 CD4+ T cells of each vaccine group over time. *p* values displayed are comparisons between the ratios outlined in black. **(d)** Representative flow charts and percentages of wildtype spike-specific memory B cells (MBCs) of all circulating B cells in blood. **(e)** The wildtype spike-specific MBC responses in the blood of each vaccine group, shown as mean and SEM over time. *p* values were determined by the Wilcoxon matched-pairs signed-rank test. **(f)** Area under the curve (AUC) of MBC population from (e). Data are shown as box-and-whiskers plots showing median, interquartile (box), and range (whiskers) or as bars with mean ± SEM (n = 6). Unless otherwise noted, the *p* values listed were determined by the unpaired Mann-Whitney test.

While no significant differences in total Th1 CD4+ T cell populations were observed between RMs immunized with Alum bolus and PNP hydrogel, examination of the polyfunctional profiles of these WT-specific CD4+ T cells revealed more nuanced findings. As quickly as the first week after priming with PNP hydrogel, more than half of the Th1 CD4+ T cells are polyfunctional (defined as IL-2+TNF-α+, IL-2+IFN-γ+, TNF-α+IFN-γ+, or IL-2+TNF-α+IFN-γ+ CD4+ T cells), whereas only around 30% of CD4+ T cells are polyfunctional following priming with Alum bolus (Figure 4c, black semi-circles indicate polyfunctionality). These differences in polyfunctionality are even greater after the boost, with over 83% of CD4+ T cells being polyfunctional with PNP hydrogel at Week 12, compared with less than 60% with Alum bolus (p = 0.015).

### PNP Hydrogel Hexapro Vaccines Extended Memory B Cell Responses in Nonhuman Primates

To assess long-term memory responses after vaccination, we measured the WT spike-specific memory B cell (MBC) population in PBMCs using spike probes as described previously (Figure 4d, S9).^62, 64, 65^ Four weeks post-prime, we observed a 2-fold higher MBC population in the RMs immunized with PNP hydrogel compared with Alum bolus (Figure 4d). Similarly, RMs vaccinated with PNP hydrogel had higher MBC populations at all time points than those vaccinated with Alum bolus over 20 weeks. Interestingly, vaccination with Alum bolus generated very little MBC population pre-boost, which only peaked post-boost (Week 12), resulting in a substantial increase from Week 4 to Week 12 (p = 0.05; Figure 4d, 4e). By contrast, a robust MBC population was reached during prime (Week 4) following PNP hydrogel immunization, at a level higher than at any time point in the Alum bolus group (p = 0.95; Figure 4d, 4e). The MBC levels were therefore similar between Week 4 and Week 12 post-PNP hydrogel vaccination (p = 0.44), suggesting that priming with PNP hydrogel could already elicit robust memory responses. Nonetheless, the MBC populations following PNP hydrogel vaccination reached a higher level post-boost, resulting in a 50% higher overall MBC AUC over time (Figure 4f).

### PNP Hydrogel Hexapro Vaccines Confer Better Immune Responses Against SARS-CoV-2 Variants of Concern

After determining that immunizing RMs with PNP hydrogel vaccines elicited more potent and durable GCs, antibody titers, CD4+ T cells, and memory responses than Alum bolus Hexapro vaccines, we sought to determine whether PNP hydrogel vaccines produced broader responses against SARS-CoV-2 variants of concern (VOC). We first assessed the binding-antibody titers over time against variants of concern-Alpha (B.1.1.7; Figure S10a), Beta (B.1.351; Figure 5a), Delta (B.1.617.2; Figure 5b), Omicron BA.1 (B.1.1.529; Figure 5c), Omicron BA.2 (Figure S10b), and Omicron BA.5 (Figure 5d). Notably, across all variants, RMs vaccinated with PNP hydrogel elicited significantly higher IgG titers as early as Week 4 (pre-boost) than those immunized with Alum bolus (p < 0.05 across all variants). Similarly, the PNP hydrogel vaccines maintained sustained and robust titers for up to 48 weeks post-immunization, resulting in significantly higher titers against all variants at Week 48 than Alum bolus vaccines (p < 0.05 across all variants).

**Figure 5.**
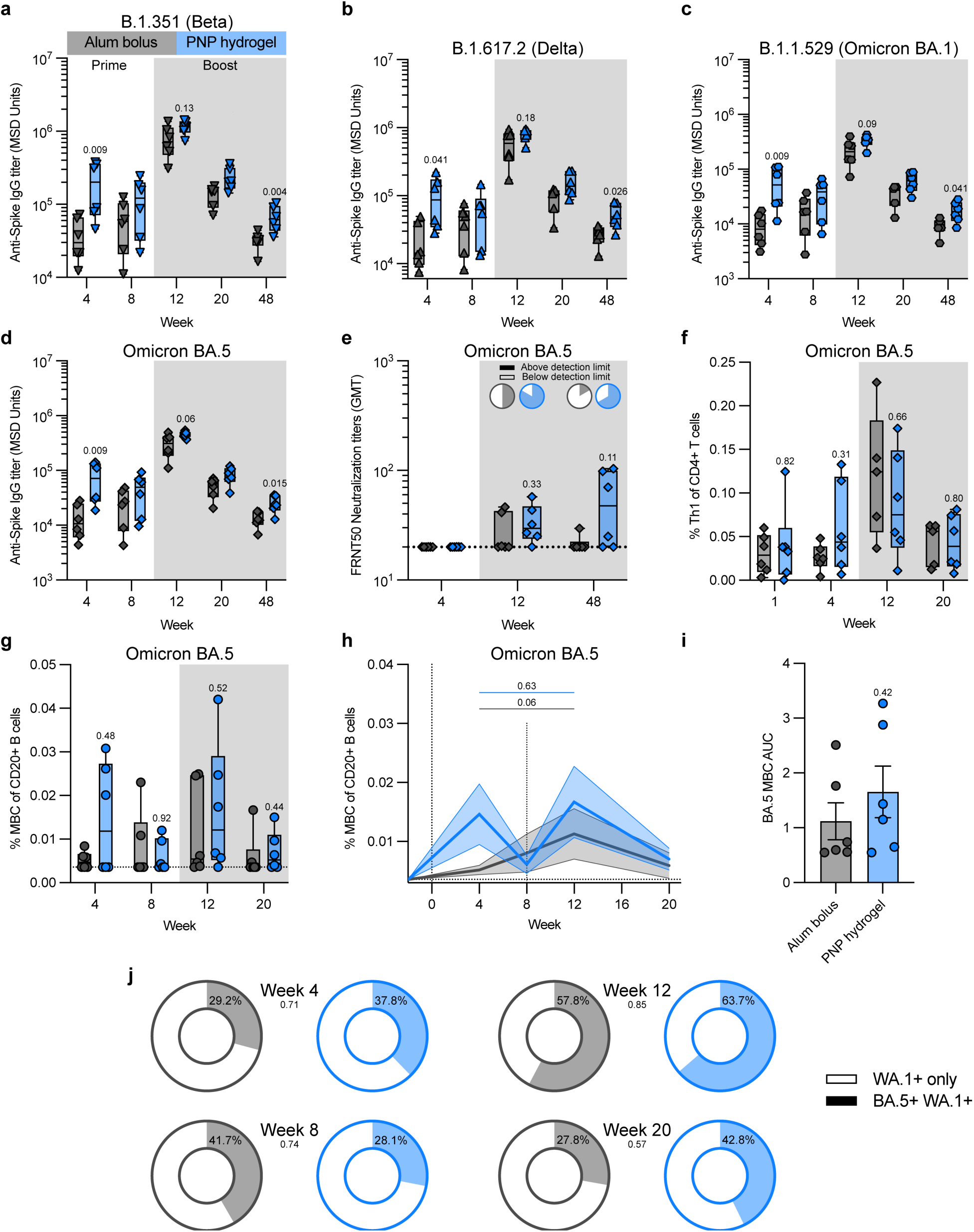
PNP Hydrogel Hexapro Vaccines Confer Better Immune Responses Against SARS-CoV-2 Variants of Concern. Anti-spike IgG titers of each vaccine group over time against the **(a)** Beta, **(b)** Delta, **(c)** Omicron BA.1, or **(d)** Omicron BA.5 variant. **(e)** Neutralizing activity of sera from immunized RMs against Omicron BA.5-type SARS-CoV-2 viruses pre- and post-boost (W4 vs W12 and W48) represented as FRNT_50_ values. Pie charts depicted the ratios of responders to nonresponders, defined as RMs with no detectable neutralizing activity. **(f)** Percentages of Omicron BA.5 spike-specific Th1 CD4+ T cells of all CD4+ T cells of each vaccine group over time. **(g)** Percentages of Omicron BA.5 spike-specific memory B cells (MBCs) of all circulating B cells in blood. **(h)** The Omicron BA.5 spike-specific MBC responses in the blood of each vaccine group, shown as mean and SEM over time. *p* values were determined by the Wilcoxon matched-pairs signed-rank test. **(i)** Area under the curve (AUC) of MBC population from (h). **(j)** Pie charts representing the fractions of wildtype only versus wildtype and Omicron BA.5 double positive MBCs of each vaccine group over time. *p* values displayed are comparisons between the double-positive MBCs. Data are shown as box-and-whiskers plots showing median, interquartile (box), and range (whiskers) or as bars with mean ± SEM (n = 6). Unless otherwise noted, the *p* values listed were determined by the unpaired Mann-Whitney test.

As time progressed and the SARS-CoV-2 virus evolved, the Omicron BA.5 variant eventually became the predominant VOC and the most difficult to neutralize due to its strong immune-evasiveness. Accordingly, we focused our subsequent studies on immune responses to the Omicron BA.5 variant. We first measured the neutralizing antibody responses against authentic SARS-CoV-2 Omicron BA.5 virus using the focus-reduction neutralization titer (FRNT) assay. We noticed that the Alum bolus vaccine resulted in only 2 out of the 6 RMs on Week 12 and 1 out of the 6 RMs on Week 48 to produce detectable neutralizing antibody titers against Omicron BA.5 (Figure 5e). Notably, the PNP hydrogel vaccine induced detectable neutralizing antibodies against Omicron BA.5 in 5 of 6 RMs at Week 12 and 4 of 6 RMs at Week 48 (p = 0.33 and p = 0.11, respectively; Figure 5e).

Analyzing Omicron-specific Th1 CD4+ T cells revealed similar findings as WT-specific CD4+ T cells. Specifically, RMs receiving PNP hydrogel vaccines generated stronger Th1 CD4+ T cells during priming than those receiving Alum bolus vaccines (Figure 5f). Consequently, immunization with PNP hydrogel vaccines led to higher Omicron BA.5-specific CD4+ T cell production over 20 weeks, as measured by AUC (Figure S11).

We then assessed memory B cell responses to Omicron BA.5 by examining Omicron BA.5-specific MBC populations. As early as Week 4 post-prime, RMs receiving PNP hydrogel vaccines generated more than double the MBC population compared to those immunized with Alum bolus vaccines, as fewer than half of the RMs elicited detectable Omicron BA.5-specific MBCs in the Alum bolus group before the boost (Figure 5g, 5h). Post-boost, PNP hydrogel vaccines elicited a second peak in the Omicron BA.5-specific MBC population and sustained a robust MBC population in four of the six RMs at Week 20 (Figure 5g, 5h). In contrast, RMs immunized with Alum bolus vaccines produced 50% less MBCs at Week 12, despite being their peak MBC level, and only two RMs had detectable Omicron BA.5-specific MBCs at Week 20. All resulted in the PNP hydrogel group generating 50% more total Omicron BA.5-specific MBCs over the 20-week period than the Alum bolus group, as measured by the AUC (1.7 versus 1.1, respectively; p = 0.42; Figure 5i).

Finally, we assessed the cross-reactivity of these MBCs against both WT and Omicron BA.5 after WT Hexapro vaccinations. Higher cross-binding MBCs at Week 4, Week 12, and Week 20 were observed after immunizing with PNP hydrogels compared with Alum bolus (Figure 5j, Figure S6). Cross-binding MBC levels waned rapidly in RMs receiving Alum bolus vaccines, with the proportion of cross-binding MBCs at Week 20 dropping to the lowest value measured at 27.8%. Encouragingly, RMs immunized with the PNP hydrogel vaccines sustained durable cross-binding MBC levels of 42.8% at Week 20, exceeding pre-boost levels.

## Discussion

Among the greatest challenges facing modern vaccinology are waning immunity and limited protection against rapidly evolving viral strains. Indeed, despite the SARS-CoV-2 pandemic spurring rapid advances in vaccine design, even the most efficacious and state-of-the-art vaccines, such as the Pfizer and Moderna mRNA vaccines and the Novavax subunit vaccine, face waning immunity against both wildtype and emerging variants of concern. For instance, protection against the Omicron variant declines to ∼45% just 10 weeks after the second mRNA booster vaccine (third immunization).^66, 67^ The constant need for booster shots against the current circulating variant to maintain protection only further exacerbates vaccine inequity, costs, and patient nonadherence. Vaccines against other infectious diseases, such as influenza and HIV, face similar challenges.

With respect to waning immunity, we show that slow delivery of Hexapro vaccines using PNP hydrogels could significantly improve both the magnitude and early persistence of vaccine responses. Immunization with PNP hydrogel resulted in significantly higher titer AUC over a 48-week period and significantly longer titer decay half-life, increased by over 3 weeks. Indeed, at W48, almost a year after vaccination, the average titer from RMs that received PNP hydrogel vaccines was more than double that from RMs that received Alum bolus vaccines at the same time point. Encouragingly, this average titer was also higher than the average titer at W8 in RMs receiving the Alum bolus vaccines. Further, the stark comparison between 4 of 6 RMs in the PNP hydrogel group and only 1 of 6 RMs in the Alum bolus group still producing detectable neutralizing titers against Omicron BA.5 48 weeks after vaccination highlights features that are consistent with improved durability of the PNP hydrogel vaccine.

In addition to enhanced magnitude, we demonstrated that PNP hydrogel vaccines significantly broadened protection, with higher binding titers against all variants tested, including the distant Omicron BA.1, BA.2, and BA.5 variants compared to Alum bolus vaccines. PNP hydrogels also led to improvements in both the magnitude and durability of neutralization titers against Omicron BA.5 compared to Alum bolus. The higher magnitude of Omicron BA.5+ Memory B cells and the higher ratio of WT+BA.5+ MBCs from the RMs immunized with PNP hydrogels would suggest improved B cell diversity, most likely resulting from increased affinity maturation from extensive GC responses. These results are consistent with our observation of more active GC responses in both mice and RMs, longer cTfh activity, and a greater proportion of polyclonal CD4+ T cells. Our findings therefore align with previous findings by Crotty et al regarding slow delivery of vaccines.^12, 13^ Previous studies in our lab have also demonstrated that PNP hydrogels could sustain and co-deliver SARS-CoV-2 spike protein, as well as other vaccine antigens such as ovalbumin, influenza hemagglutinin, the SARS-CoV-2 receptor-binding domain proteins and nanoparticles, and HIV MD39 proteins, with small-molecule adjuvants such as the TLR7/8a adjuvant 3M-052 for a tunable period of 2-4 weeks.^30–41^ In this study, we’ve shown that, by doing so with Hexapro and 3M-052, we’ve improved immune memory responses over time, including more diverse B cell and polyclonal CD4+ T cell responses than in the Alum bolus control, leading to improved magnitude, early persistence, and breadth of the humoral responses.

Separately, robust priming responses after immunization with PNP hydrogels could suggest its use as a single-shot vaccine platform. We observed that all mice receiving the hydrogel vaccine, but not all mice receiving the Alum bolus vaccine, seroconverted after just the priming dose. Additionally, mice primed with hydrogel vaccines elicited higher average titers during prime than mice vaccinated with Alum bolus did at any time, even post-boost. Similarly, titers during prime from RMs vaccinated with PNP hydrogels were higher or equivalent to the W20 titers (12W after boost) from RMs vaccinated with Alum bolus. Furthermore, one of the biggest challenges in vaccine priming alone is the inability to elicit a robust, long-lasting memory response. Indeed, most RMs vaccinated with Alum bolus did not generate any MBCs against Omicron BA.5 after priming (only 2 of 6). In contrast, a strong peak in the MBC population (against both WT and Omicron BA.5) was observed rapidly after RMs were immunized with PNP hydrogel. Similar to previous studies,^36, 40^ this shows great promise for using PNP hydrogels to reduce vaccine regimens and costs, increase patient compliance, and improve the efficacy of one-dose vaccines.

In summary, we report a biomaterial approach using an injectable PNP hydrogel depot technology for the slow delivery of Hexapro vaccines in Rhesus macaques. We demonstrated that PNP hydrogel vaccines improved vaccine priming responses, increased the magnitude and persistence of antibody binding titers, enhanced germinal center responses, enriched Th1 CD4+ T cell and memory B cell responses, and broadened responses against variants of concern. Together, as the first reported study of a biomaterial for sustained vaccine delivery in NHP, we established that PNP hydrogel vaccines are promising clinical candidates for slow-release vaccination to enhance humoral responses.

## Materials and Methods

### Materials

Poly(ethylene glycol)methyl ether 5000 Da (PEG), 3,6-Dimethyl-1,4-dioxane-2,5-dione (Lactide), 1,8-diazabicyclo(5.4.0)undec-7-ene (DBU, 98%), (Hydroxypropyl)methyl cellulose (HPMC, meets USP testing specifications), N,N-Diisopropylethylamine (Hunig’s base), N-methyl-2-pyrrolidone (NMP), 1-dodecyl isocyanate (99%), mini Quick Spin Oligo columns (Sephadex G-25 Superfine packing material), Sepharose CL-6B crosslinked, bovine serum albumin (BSA), Acetonitrile (ACN), Dimethyl Sulfoxide (DMSO), and dichloromethane (DCM) were purchased from Sigma-Aldrich. Hexapro was kindly provided by the Institute for Protein Design at University of Washington. 3M-052 and 3M-052/Alum (AAHI-AL030) were purchased from 3M and the Access to Advanced Health Institute (AAHI). Unless otherwise stated, all chemicals were used as received without further purification.

### Preparation of HPMC-C_12_

HPMC-C_12_ was prepared according to a previously reported procedure.^30–41^ Briefly, hypromellose (HPMC, 1.5 g) was dissolved in 60 mL of anhydrous NMP. The solution was then heated at 50 °C for 30 mins. A solution of dodecyl isocyanate (0.75 mmol, 183 μL) in 5 mL of anhydrous NMP was added dropwise, followed by 105 μL of *N*,*N*-diisopropylethylamine (0.06 mmol). The solution was stirred at room temperature for 20 h. The polymer was recovered from precipitation in acetone and filtered. The polymer was purified through dialysis (3 kDa mesh) in Milli-Q water for 4 days and lyophilized to yield a white amorphous polymer. The polymer mixture was then lyophilized and reconstituted to a 60 mg/mL solution in sterile PBS 1X.

### Preparation of PEG-PLA NPs

PEG-*b*-PLA was prepared as previously reported.^30–41^ Prior to use, commercial lactide was recrystallized in ethyl acetate and DCM were dried via cryo distillation. PEG-methyl ether (5 kDa, 0.25 g, 4.1 mmol) and DBU (15 µL, 0.1 mmol) were dissolved in 1 mL of dry DCM under nitrogen atmosphere. Lactide (1.0 g, 6.9 mmol) was dissolved in 4.5 mL of dry DCM under nitrogen atmosphere. The lactide solution was then quickly added to the PEG/DBU mixture and was allowed to polymerize for 8 min at room temperature. The reaction was then quenched with an acetic acid solution and the polymer precipitated into a 1:1 mixture of ethyl ether and hexanes, collected by centrifugation, and dried under vacuum. NMR spectroscopic data, M_n_, and dispersity were then confirmed to match those previously described.

PEG-*b*-PLA NPs were prepared as previously described. A 1 mL solution of PEG-b-PLA in 75:25 ACN:DMSO (50 mg/mL) was added dropwise to 10 mL of Milli-Q water stirring at 600 rpm. The hydrodynamic diameter of the NPs was measured on a DynaPro II plate reader (Wyatt Technology). Three independent measurements were performed for each sample. The particle solution was purified using centrifugal filters (Amicon Ultra, MWCO 10 kDa) at 4500 RCF for 1 h and resuspended in 1X PBS to a final concentration of 200 mg/mL.

### Preparation of Hexapro

Hexapro was synthesized, purified, and tested for antigenicity as previously described.^68, 69^ Briefly, The Hexapro protein was synthesized and purified using recombinant DNA and mammalian cell expression methods. A plasmid encoding Hexapro with a secretion signal and C-terminal histidine tag was constructed and amplified in *E. coli* (New England Biolabs C2987H-81), then purified using a DNA MaxiPrep system (Macherey-Nagel DNA MaxiPrep kit). This plasmid was transfected into Expi293F mammalian cells (Thermo Fisher Α14527) grown in suspension culture, using polyethylamine to facilitate uptake. After 6 days of incubation at 33 °C under 8% CO2 with constant agitation at 150 RPM, the cells expressed the Hexapro protein, which was collected from the culture supernatant. The protein was purified through immobilized metal affinity chromatography (IMAC) using nickel resin, followed by concentration and dialysis into a stabilized buffer.

The purified protein was characterized for concentration, purity, and endotoxin levels. Spectrophotometric analysis using the Stunner Quantification system measured absorbance across wavelengths to calculate protein concentration via Beer’s Law. Endotoxin contamination was assessed using a Limulus Amebocyte Lysate (LAL) assay. Protein purity and structural integrity were evaluated through SDS-PAGE electrophoresis under both reduced and non-reduced conditions, allowing visualization of Hexapro bands and confirmation of expected molecular weight.

Functional and structural analyses were also performed to assess antigenicity and stability. Binding interactions between Hexapro and known receptors or antibodies (hACE2-Fc, LyCov555, and B38) were measured using biolayer interferometry on a ForteBio Octet system, tracking association and dissociation kinetics. Additionally, samples both before and after a freeze–thaw cycle were examined to evaluate stability. Structural visualization was performed using negative-stain electron microscopy, where stained protein samples on carbon grids were imaged at high magnification and processed with CryoSPARC software to generate two-dimensional class averages of the protein particles.

### Preparation and Formulations of PNP Hydrogels

Polymer-nanoparticle (PNP) hydrogels were formed with HPMC-C_12_ and a mixture of PEG-*b*-PLA NPs in TBS 1X. PNP-2-10 (2 wt% HPMC-C_12_ and 10 wt% NPs) hydrogels were prepared by mixing one-third of the final volume with 6 wt% HPMC-C_12_ solution, half of the final volume with 20 wt% NPs solution, and the remaining volume (one-sixth) with TBS 1X to achieve the PNP-2-10 formulation described above. PNP-1-5 (1 wt% HPMC-C_12_ and 5 wt% NPs) hydrogels were prepared by mixing one-sixth of the final volume with 6 wt% HPMC-C_12_ solution, a quarter of the final volume with 20 wt% NPs solution, and the remaining volume (7/12) with TBS 1X to achieve the PNP-1-5 formulation described above. Based on the desired antigen and adjuvant formulations, aqueous adjuvant (3M-052) and antigen (Hexapro) were included as part of the PEG-b-PLA NPs and 1X TBS mixture by subtracting the volume of the antigen and adjuvant cargoes from the volume of 1X TBS. The hydrogels were formed by mixing the solutions using syringes connected through an elbow mixer as previously reported.^30–41^

### Hydrogel Rheological Characterization

Rheological characterization was completed on a Discovery HR-2 Rheometer (TA Instruments). Measurements were performed using a 20 mm serrated-plate geometry at 25 °C and a 500 µm gap height. Empty PNP-2-10 or vaccine-loaded (Hexapro and 3M-052) PNP-2-10 hydrogels were loaded on the geometry. Dynamic oscillatory frequency sweeps were conducted at a constant strain of 1% and angular frequencies from 0.1 to 100 rad/s. Amplitude sweeps were performed at a constant angular frequency of 10 rad/s from 0.5% to 10,000% strain. Flow sweep and steady shear experiments were performed at shear rates from 50 to 0.005 1/s, whereas stress-controlled flow sweep measurements were conducted at shear rates from 0.001 to 10 1/s. Step-shear experiments were performed by alternating between low shear rates (0.1 rad/s for 60 s) and high shear rates (10 rad/s for 30 s) for three full cycles. Yield stress values were extrapolated from stress-controlled flow sweep and amplitude sweep measurements.

### Hydrogel Extrudability

The extrudabilities of both empty and loaded PNP hydrogels were modeled using the theoretical framework established by Lopez Hernandez et al.^48, 53^ Briefly, the consistency index (K) and shear-thinning index (n) were determined by fitting a power-law regression to the flow rheology data in Excel. These fitted parameters were then input into the Lopez Hernandez model to generate an Ashby-style plot using the following process constraints: maximum pressure = 26 kPa; minimum flow rate = 6 mL min−1; needle inner radius = 152 microns (21G needle); needle length = 24.5 mm.

### Hydrogel Injection Force Characterization

Force of injection was quantified by measuring the force required to inject a hydrogel through a known needle gauge at a known flow rate using a syringe of known barrel dimensions.^54^ A force sensor was built that encompassed a load cell [FUTEK LLB300 50 lb Subminiature Load Button (model no. LLB300, item no. FSH03954, serial no. 705242)] attached to a syringe pump [KD Scientific Syringe Pump (model no. LEGATO 100, catalog no. 788100, serial no. D103954)]. An Omega Engineering Platinum Series Meter (model no. DP8PT, serial no. 18110196) was used to convert load-cell resistance measurements to force in kilograms (kg). The load cell was calibrated before measuring the injection force.

Injection force experiments were performed as follows. A 1-mL Luer-lock syringe (Air-Tite Products Co.) containing either PNP-2-10 or PNP-1-5 hydrogels, fitted with a 21G, 1-inch blunt-tip needle, was loaded into the syringe pump. The syringe pump height was adjusted to bring the force sensor load button into contact with the end of the syringe plunger. The initial force was at or near 0 kg. The appropriate syringe barrel dimensions as well as the desired flow rate and injection volume were then selected. The syringe pump moved at the programmed rate, injecting the hydrogels through the attached needle. The force sensor, coupled to the Omega unit, measured the force required to inject the protein suspension at the desired flow rate. A LabVIEW program recorded forces throughout an injection experiment and displayed a graph of injection force over time. Force of injection was quantified by subtracting the average initial force (background) from the average plateau injection force. The injection force in kilograms was converted to newtons by multiplying by 9.81.

### Vaccine Formulations for Maurine Study

The vaccines contained 0.5 μg of Hexapro and 1 μg of 3M-052 adjuvant per dose, either in bolus form or in PNP hydrogels. For bolus groups, 3M-052/Alum (1 μg + 100 μg, respectively) was combined with Hexapro in TBS 1X to a volume of 100 μL per dose and loaded into syringes with a 26-gauge needle for subcutaneous injection. Mice were boosted in Week 8. Hydrogel vaccines were prepared as described above for the PNP-2-10 hydrogel, to a volume of 100 μL per dose, in syringes fitted with a 21-gauge needle for subcutaneous injection.

### Mice and Vaccination

Six-to-seven-week-old female C57BL/6 (B6) mice were purchased from Charles River and housed in the animal facility at Stanford University. Mice were shaved to receive a subcutaneous injection of 100 µL of soluble or hydrogel vaccine on the right side of their flank under brief anesthesia. Mouse blood was collected weekly from the tail veins.

### Mouse Serum Enzyme-Linked Immunosorbent Assay (ELISAs)

Serum antigen-specific IgG antibody endpoint titers were measured using an endpoint ELISA. MaxiSorp plates (Invitrogen) were coated with SARS-CoV-2 spike protein (Sino Biological 40589-V08H4) at 2 µg/mL in PBS 1X overnight at 4 °C and subsequently blocked with PBS 1X containing 1 wt% BSA for 1 h at 25 °C. Plates were washed 5 times between steps in 1X PBS containing 0.05 wt% Tween-20. Serum samples were diluted in diluent buffer (PBS 1X with 1 wt% BSA) to 1:100, then serially diluted 4-fold, and incubated in the previously coated plates for 2 h at 25 °C. Goat-anti-mouse IgG Fc-HRP (1:10,000, Invitrogen A16084) was then added for 1 h at 25 °C. The plates were developed with TMB substrate (high-sensitivity TMB from Abcam), and the reaction was stopped with 1 M HCl. Absorbance was measured using a Synergy H1 microplate reader (BioTek Instruments) at 450 nm. Data were analyzed in GraphPad Prism and fit with a five-point asymmetric sigmoidal curve with constraints S > 0, top < 4 (maximum limit of detection), and bottom = 0.03 (background absorbance value). The endpoint titer was defined as the serum dilution at which the absorbance was 2X the background value (0.1), interpolated from the curve fits in GraphPad Prism.

### Pseudotyped Lentivirus Production and Mice Serum Neutralization Assays

SARS-CoV-2 spike-pseudotyped lentiviruses encoding a luciferase-ZsGreen reporter were produced in HEK293F cells by co-transfection of five plasmids. The five plasmids include a packaging vector (pHAGE-Luc2-IRES-ZsGreen), a plasmid encoding the SARS-CoV-2 spike (HDM-SARS2-spike-delta21, Addgene, 155130), and three helper plasmids (pHDM-Hgpm2, pHDM-Tat1b, and pRC-CMV_Rev1b). 50 mL of cells were diluted to a density of approximately 3–4 × 10^6^ cells per mL. The transfection mixture was prepared by adding five plasmids (50 µg of packaging vector, 17 µg of SARS-CoV-2-encoding plasmid, and 11 µg of each helper plasmid) to 5 ml of Expi-free medium, followed by the dropwise addition of BioT transfection reagent (150 µl, Bioland Scientific) with vigorous mixing. After 10 min of incubation at room temperature, the transfection mixture was transferred to HEK293F cells. D-glucose (4 g/L, Sigma-Aldrich) and valproic acid (3 mM, Acros Organics) were then added to the cells immediately post-transfection to increase recombinant protein production. The cells were harvested 3 days after transfection by spinning the cultures at 300 *g* for 5 minutes. The supernatant was filtered through a 0.45-µm filter and 0.5 mL of 1 mM HEPES was added to neutralize the pH. Viral stocks were aliquoted and flash-frozen in liquid nitrogen. They were stored at −80 °C and titrated before further use.

Antisera were heat-inactivated (56 °C, 30–60 min) before neutralization assays. Neutralization against SARS-CoV-2 WT (Wuhan) was analyzed in HeLA-ACE2/TMPRSS2 cells. One day before infection (day 0), cells were seeded at 8,000 cells per well in white-walled, white-bottom, 96-well plates (Thermo Fisher or Greiner Bio-One). On day 1, antisera were serially diluted in D10 media and mixed 1:1 with pseudoviruses for 1-2 hrs at 37 °C before being transferred to cells. The pseudovirus mixture contained SARS-CoV-2 WT, D10 media, and polybrene (1:500). Assays were read out with luciferase substrates 2 d after infection by removing the media from the wells and adding 80 uL of a 1:1 dilution of BriteLite in DPBS (BriteLite Plus, Perkin Elmer). Luminescence values were measured using a microplate reader (BioTek Synergy^™^ HT or Tecan M200). NT_50_ was calculated as the serum dilution where 50% inhibition of infection was observed. Percent infection was normalized on each plate by averaging cell-only (0% infectivity) and virus-only (100% infectivity) wells. Neutralization assays were performed in technical duplicates.

### Murine Flow Cytometry of Draining Lymph Nodes

Mice were euthanized using carbon dioxide at different time points post-immunization, as shown in Figure 2, to collect the inguinal lymph nodes. Following their dissociation into single-cell suspensions, cells were stained for viability with Ghost Dye Violet 510 (Tonbo Biosciences, cat. number 13-0870-T100) for 5 min on ice, then washed with FACS buffer (PBS 1X with 2% FBS, 1 mM EDTA). Then, Fc receptors were blocked using anti-CD16/CD38 antibody (clone: 2.4G2, BD Biosciences, cat. Number 553142) for 5 mins on ice and stained with fluorochrome conjugated antibodies: CD19 (PerCP-Cy5.5, clone: 1D3, BioLegend, cat. Number 152406), CD95 (PE-Cy7, clone: Jo2, BD Biosciences, cat. Number 557653), GL7 (AF488, clone: GL7, BioLegend, cat. Number 144613), CD3 (AF700, clone: 17A2, BioLegend, cat. Number 100216), CD4 (BV650, clone: GK1.5, BioLegend, cat. Number 100469), CXCR5 (BV711, clone: L138D7, BioLegend, cat. Number 145529), and PD1 (PE-Dazzle^TM^594, clone: 29F.1A12, BioLegend, cat. Number 135228) for 30 mins on ice. Cells were washed and fixed with 4% PFA on ice. After washing, cells were analyzed using a Symphony II flow cytometer (BD Biosciences). Data were analyzed with FlowJo 10 (FlowJo, LLC).

### Enzyme-Linked Immunosorbent Spot (ELISpot)

Single-color mouse IgG ELISPOT kits were purchased from Cellular Technology Limited. Bone marrow cells were harvested following immunization schedules outlined in Figure 2. Wells were first coated with 80 µL of Hexapro at 25 µg/mL in PBS 1X or Capture Solution (for positive control to detect all antibody-secreting cells) prepared according to the manufacturer’s specifications overnight at 4 °C and subsequently blocked with assay medium (RPMI 1640 with 10% FBS, 2mM L-glutamine, 100 U/ml Penicillin, 100 μg/mL Streptomycin, 8 mM HEPES, and 50 μM 2-mercaptoethanol). 300,000 bone marrow cells or spleen cells per sample were pipetted into the wells and incubated for 5 hours. Spots were then developed following the manufacturer’s instructions (Immunospot S6 Ultra M2).

### Vaccine Formulations for Nonhuman Primate Study

The vaccines contained 5 μg of Hexapro and 12.5 μg of 3M-052 adjuvant per dose, either as a bolus or in PNP hydrogels. For bolus groups, 3M-052/Alum (5 μg + 500 μg, respectively) was combined with Hexapro in TBS 1X to a volume of 500 μL per dose and loaded into syringes with a 26-gauge needle for injection. Hydrogel vaccines were prepared as described above for the PNP-1-5 hydrogel, to a volume of 250 μL per dose, in syringes fitted with a 21-gauge needle for injection. NHPs were subcutaneously injected at the right deltoid and boosted in Week 8.

### Rhesus Macaques Animal Subjects and Vaccinations

Twelve male rhesus macaques of Indian origin, aged 5 to 15 years were purchased from the Tulane National Primate Research Center and housed and maintained as per National Institutes of Health (NIH) guidelines at the New Iberia Research Center facilities, University of Louisiana at Lafayette in accordance with the rules and regulations of the Committee on the Care and Use of Laboratory Animal Resources. The animals were fed a nonhuman primate diet (Teklad) supplemented regularly with fresh fruits or produce; water was provided ad libitum. All animals were negative for simian immunodeficiency virus, simian T cell leukemia virus, and simian retrovirus. The entire study was reviewed and approved by the University of Louisiana at Lafayette Institutional Animal Care and Use Committee (IACUC protocol number: 2022-007-8832-NIRC). All animal collections were done under sedation.

### NHP Fine Needle Aspirate of Draining Lymph Nodes and Germinal Center Flow Cytometry

GC Tfh cells and activated B cells were characterized by flow cytometry of fresh lymph node cells obtained via fine-needle aspirates. 0.5 to 1 million cells were stained as follows. Initially, cells were washed in PBS. Cells were incubated with 100 µL of fixable viability dye eFlour506 (Thermo Fisher, cat. no. 65-0866-14) diluted 1:1000 in PBS for 20 min at room temperature. After washing with 2mL of PBS-5% FCS, cell surfaces were stained at RT for 25 min in 100 µL of an antibodies cocktail including: CD4 BV785 (BioLegend, clone OKT4), CD3 APC-Cy7 (BD, clone SP34-2), CD20 PE-Cy7 (BioLegend clone 2H7), CXCR5 PE (eBiosciences, clone MU5UBEE), PD-1 BV421 (BioLegend, clone EH12.2H7) and Brilliant Buffer Stain Plus (BD). Following two washes with 2 mL of PBS-5% FCS, cells were incubated 20 min at room temperature with 150 µL of fix/perm buffer (eBiosciences Foxp3/transcription buffer set Cat#00-5523-00). Cells were then washed with 2ml of Perm/Wash buffer and stained for 25 min at room temperature with antibodies: Ki-67-FITC (BD, clone B56) and Bcl-6-AF647 (BD, clone K112-91), each diluted to 100 µL in Perm/Wash buffer. Cells were washed twice with 2 mL of Perm/Wash and fixed for 10min at room temperature in 150 µL of BD Cytofix. Finally, cells were washed with 2 mL of PBS-5 % FCS and suspended in 250 µL of PBS-5 % FCS. Cell data were acquired on a BD FACS Aria Fusion cytometer, and analysis was performed using FlowJo software. Graphing and statistical analysis were done using Prism software.

### Anti–Spike Protein Electrochemiluminescence Binding Enzyme-Linked Immunosorbent Assay

Anti–Spike protein IgG titers were measured using V-PLEX SARS-CoV-2 Key Variant Spike Panel 1 (IgG) Kit from Meso Scale Discovery (MSD; catalog no. K15651U). The assay was performed as per the manufacturer’s instructions. Briefly, the multispot 96-well plates were blocked in 0.15 mL of blocking solution with shaking at 700 rpm at room temperature. After 30 min of incubation, 50 μL of serum was diluted in antibody diluent, and serially diluted calibrator solutions were added to each plate in the designated wells. The plates were incubated at room temperature with shaking for 2 hours. After 2 hours of incubation, the plates were washed, and 50 μL of Sulfo-tag–conjugated anti-IgG was added, and the plates were incubated at room temperature for 1 hour. After incubation, the plates were washed, and 0.15 mL of MSD GOLD Read buffer was added. The plates were immediately read using the MSD instrument. The unknown concentrations were extrapolated using a standard curve drawn using the calibrators in each plate and presented as relative MSD AU/mL.

### Antibody Half-Life Model

We followed previously reported methods to measure antibody decay.^36^ Briefly, parametric bootstrapping method was used in Matlab by assuming the underlying distribution of each time point was to be log-normally distributed. For each of the 1000 simulations in the bootstrap, linear regression was used to fit the sample to an exponential decay model, yielding a distribution of half-lives.

### Viruses and cells for focus reduction neutralization test

VeroE6-TMPRSS2 cells were described previously and cultured in complete DMEM (DMEM with 10% FBS + penicillin-streptomycin) in the presence of Gibco puromycin (10 mg/mL; #A11138-03). nCoV/USA_WA1/2020 (WA/1) was propagated from an infectious SARS-CoV-2 clone as previously described.^70^ icSARS-CoV-2 was passaged once to generate a working stock. The hCoV-19/USA/MD-HP01542/2021 (B.1.351) was provided by A. Pekosz (Johns Hopkins University) and was propagated in Vero-TMPRSS2 cells.^71^ hCoV19/EHC_C19_2811C (referred to as the B.1.1.529 variant) was derived from a mid-turbinate nasal swab collected in December 2021. This SARS-CoV-2 genome is available under Global Initiative on Sharing Avian Influenza Data accession number EPI_ISL_7171744. All viruses used in the focus reduction neutralization test (FRNT) assay were deep-sequenced and confirmed as previously described.^72^

FRNT assays were performed as previously described.^72–74^ Briefly, samples were diluted threefold in eight serial dilutions in DMEM, in duplicate, with an initial dilution of 1:10, for a total volume of 60 μL. Serially diluted samples were incubated with an equal volume of WA1, BA.5 (100-200 foci per well, depending on the target cell density) at 37 °C for 45 min in a round-bottomed 96-well culture plate. The antibody-virus mixture was then added to VeroE6-TMPRSS2 cells and incubated at 37° C for 1 hour. After incubation, the antibody-virus mixture was removed, and 100 μl of prewarmed 0.85% methylcellulose (Sigma-Aldrich, #M0512-250G) overlay was added to each well. Plates were incubated at 37°C for either 18 hours (WA1 or B.1.351) or 40 hours (B.1.1.529), and the methylcellulose overlay was removed and washed six times with PBS. Cells were fixed with 2% paraformaldehyde in PBS for 30 min. After fixation, plates were washed twice with PBS, and permeabilization buffer (0.1% bovine serum albumin and 0.1% saponin in PBS) was added to permeabilized cells for at least 20 min. Cells were incubated with an anti–SARS-CoV Spike protein primary antibody directly conjugated to Alexa Fluor 647 (CR3022-AF647) for 4 hrs at room temperature or overnight at 4 °C. Cells were washed three times in PBS, and foci were visualized on an ELISpot reader. Antibody neutralization was quantified by counting the number of foci for each sample using the Viridot program.^75^ The neutralization titers were calculated as follows: 1 − (ratio of the mean number of foci in the presence of serum and foci at the highest dilution of the respective serum sample). Each specimen was tested in duplicate. The FRNT-50 titers were interpolated using a four-parameter nonlinear regression in GraphPad Prism 9.2.0. Samples that did not neutralize at the limit of detection at 50% were plotted at 10 and used for geometric mean and fold change calculations.

### T cell ICS assay

Antigen-specific T cell responses were measured using the ICS assay. Cryopreserved PBMCs were revived, counted, and resuspended at a density of 2 × 10^6^ live cells/mL in complete RPMI 1640 (RPMI 1640 supplemented with 10% FBS and penicillin-streptomycin). The cells were rested overnight at 37 °C in a CO_2_ incubator. The next morning, the cells were counted again, resuspended at a density of 12 × 10^6^/mL in complete RPMI 1640, and 100 μL of the cell suspension containing 1.2 × 10^6^ cells was added to each well of a 96-well round-bottomed tissue culture plate. Each sample was treated with three or four conditions depending on cell numbers: no stimulation or a peptide pool spanning the Spike protein of the ancestral Wuhan strain or Omicron BA.5 variant (where cell numbers permitted) in the presence of anti-CD28 (1 μg mL^−1^; clone CD28.2, BD Biosciences) and anti-CD49d (clone 9F10, BD Biosciences), as well as anti-CXCR3 and anti-CXCR5. The peptide pools consisted of 15-nucleotide oligomers with 10-nucleotide overlaps spanning the entire Spike protein sequence of each variant. Each peptide was dissolved at a concentration of 20 mg/mL in dimethyl sulfoxide (DMSO), and individual peptides were pooled to prepare each variant-specific peptide pool after sequential lyophilization, as previously reported.^7^ PBMCs were stimulated at a final concentration of 1 μg/mL of each peptide in the final reaction with an equimolar amount of DMSO [0.5% (v/v) in 0.2-mL total reaction volume] as a negative control. The samples were incubated at 37 °C in CO_2_ incubators for 2 hours before the addition of brefeldin A (10 μg mL^−1^). The cells were incubated for an additional 4 hrs. The cells were washed with PBS and stained with Zombie ultraviolet (UV) fixable viability dye (BioLegend). The cells were washed with PBS containing 5% FBS prior to the addition of the surface antibody cocktail (Table S1). The cells were stained for 20 min at 4°C in 100 μL of staining buffer. Subsequently, the cells were washed, fixed, and permeabilized with Cytofix/Cytoperm buffer (BD Biosciences, #555028) for 20 min. The permeabilized cells were stained with ICS antibodies for 20 min at room temperature in 1× Perm/Wash buffer (BD Biosciences, #555028). The details of antibodies used in the assay are provided in Table S1. Cells were then washed twice with Perm/Wash buffer and once with staining buffer before acquisition using the BD FACSymphony flow cytometer and the associated BD FACSDiva software. All flow cytometry data were analyzed using FlowJo software v10 (TreeStar Inc.).

### Spike Protein–specific Memory B Cell Staining

Cryopreserved PBMCs were thawed and washed twice with 10 mL of fluorescence-activated cell sorting (FACS) buffer (1× PBS containing 2% FBS and 1 mM EDTA) and resuspended in 100 μl of PBS containing Zombie UV LIVE/DEAD dye at 1:200 dilution (BioLegend, #423108) and incubated at room temperature for 15 min. After washing, cells were incubated with an antibody cocktail for 1 hour on ice, protected from light. The following antibodies were used: IgD phycoerythrin (PE; Southern Biotech, #2030-09), IgM peridinin chlorophyll protein–Cy5.5 (BioLegend, #314512), CD20 allophycocyanin-H7 (BD Biosciences, #560734), CD27 PE-Cy7 (BioLegend, #302838), CD14 brilliant violet (BV) 650 (BioLegend, #301836), CD16 BV650 (BioLegend, #302042), IgG brilliant UV 395 (BD Biosciences, #564229), CD3 BV650 (BD Biosciences, #563916), CD21 PE-CF594 (BD Biosciences, #563474), Alexa Fluor 647–and BV421 double labeled Wuhan Spike protein (Sino Biological, # 40589-V49H-B), and Alexa Fluor 488–and BV711 double labeled BA.5 Spike protein (Sino Biological, # 40589-V49H5-B-20). All antibodies were used as per the manufacturer’s instructions, and the final concentration of each probe was 0.1 μg/ml. Cells were washed twice in FACS buffer and immediately acquired on a BD FACSAria III. FlowJo software v10 (TreeStar Inc.) was used for data analysis.

### Animal Protocol

Mice were cared for following the Institutional Animal Care and Use guidelines. All animal studies were performed in accordance with the National Institutes of Health guidelines and the approval of the Stanford Administrative Panel on Laboratory Animal Care (protocol APLAC-32109). The entire NHP study was reviewed and approved by the University of Louisiana at Lafayette Institutional Animal Care and Use Committee (IACUC protocol number: 2022-007-8832-NIRC) and Stanford Administrative Panel on Laboratory Animal Care (protocol APLAC-34408). All NHP animal collections were done under sedation.

### Statistical Analysis

For *in vivo* experiments, animals were caged or treatment blocked. Mead’s Resource Equation was used to calculate the sample size above which additional subjects (n) will not have a significant impact on power. For most experiments, a sample size of *n* = 6 per group was used. No animals were considered outliers. Data are presented as mean ± standard error of mean (SEM) or median ± interquartile range as specified in the figure captions. Comparison between two groups (Alum bolus versus PNP hydrogel) at a time point was assessed using a two-tailed nonparametric Mann-Whitney unpaired rank-sum test. Data were used in their raw form or transformed (log10) as needed to maximize the coefficient of determination and improve model fit. Statistical significance was considered as *p* < 0.05. All statistical analyses were performed using GraphPad Prism.

## Data Availability

The data that support the findings of this study are available from the corresponding author upon reasonable request.

## Supporting Information

The following files are available free of charge.

Additional tables and figures of synthesis, purification, and characterization of PNP hydrogels and additional immune response data and figures. (PDF)

## Author Contributions

Conceptualization: BSO, MH, BP, EAA.

Methodology: BSO, MH, AU, NE, AS, FV, BP, EAA

Investigation: BSO, MH, JY, JMS, OMS, VL, YES, AG, JHK, AS, AU, MSS, NE, YF, JB, KAR, LMS, GJA, ASV, RR, BP, EAA

Visualization: BSO

Supervision: NPK, JF, FV, BP, EAA Resources: NPK, BP, EAA

Validation: BSO, MH, AU, KAR, NPK, JF, FV, BP, EAA

Formal analysis: BSO, MP, AU, KAR Writing—original draft: BSO, MH

Writing—review & editing: BSO, MH, JY, OMS, YES, JHK, JB, BP, EAA

All authors have given approval to the final version of the manuscript. ‡ These authors contributed equally to this work.

## Conflict of Interest Statement

EAA is listed as an inventor on several issued and pending patents related to the PNP hydrogel materials described in this manuscript. EAA is an equity holder and advisor to Appel Sauce Studios LLC, which holds an exclusive license to several patents related to the PNP hydrogel materials discussed in this manuscript. BP serves or has served on the Scientific Advisory Board of Institut Pasteur, Pfizer Vaccines, Candel Therapeutics, IMU Biosciences, Pharmajet, Techimmune, GSK, Sanofi, Medicago, Boehringer Ingelheim, Pharmajet, Icosavax, and Ed-Jen. All other authors have no conflicts to declare.

## Acknowledgment

The authors would like to thank the members of the Appel and Pulendran lab and their collaborators. We thank everyone at the NIRC for their work and care for the NHPs. The authors thank Jodi Hanson for her consultations on ELISpot protocols. Flow cytometry data were collected on an instrument at the Stanford Shared FACS Facility using an NIH S10 Shared Instrument Grant (1S10OD026831-01). We acknowledge Andrew Borst and Zhitong Peng for their contributions to negative-stain electron microscopy, which enabled us to characterize Hexapro. This work was financially supported by the Bill & Melinda Gates Foundation (INV-027411). BSO is grateful for an Eastman Kodak Fellowship. JY, OMS, and NE were supported by the National Science Foundation Graduate Research Fellowship. OMS was also supported by the Hancock Fellowship of the Stanford Graduate Fellowship in Science and Engineering. JB is grateful for a Marie-Curie fellowship from the European Union (H2020, No. 101030481). AU was supported by Stanford University Medical Scientist Training Program grants T32-GM007365 and T32GM145402. ASV, RR, and NPK are grateful of National Institute of Allergy and Infectious Diseases grant P01AI167966.

## References

1. Mascola, J.R. & Fauci, A.S. Novel vaccine technologies for the 21st century. Nature Reviews Immunology 20, 87–88 (2020).

2. Feikin, D.R. et al. Duration of effectiveness of vaccines against SARS-CoV-2 infection and COVID-19 disease: results of a systematic review and meta-regression. Lancet 399, 924–944 (2022).

3. Goldberg, Y. et al. Waning Immunity after the BNT162b2 Vaccine in Israel. N Engl J Med 385, e85 (2021).

4. Hoffmann, M. et al. The Omicron variant is highly resistant against antibody-mediated neutralization: Implications for control of the COVID-19 pandemic. Cell 185, 447–456 e411 (2022).

5. Dejnirattisai, W. et al. SARS-CoV-2 Omicron-B.1.1.529 leads to widespread escape from neutralizing antibody responses. Cell 185, 467–484 e415 (2022).

6. Cameroni, E. et al. Broadly neutralizing antibodies overcome SARS-CoV-2 Omicron antigenic shift. Nature 602, 664–670 (2022).

7. Tarke, A. et al. SARS-CoV-2 vaccination induces immunological T cell memory able to cross-recognize variants from Alpha to Omicron. Cell 185, 847–859 e811 (2022).

8. Lee, J.H. & Crotty, S. HIV vaccinology: 2021 update. Semin Immunol 51, 101470 (2021).

9. Havenar-Daughton, C. et al. Direct Probing of Germinal Center Responses Reveals Immunological Features and Bottlenecks for Neutralizing Antibody Responses to HIV Env Trimer. Cell Reports 17, 2195–2209 (2016).

10. Cirelli, K.M. & Crotty, S. Germinal center enhancement by extended antigen availability. Curr Opin Immunol 47, 64–69 (2017).

11. Crotty, S. T Follicular Helper Cell Biology: A Decade of Discovery and Diseases. Immunity 50, 1132–1148 (2019).

12. Cirelli, K.M. et al. Slow Delivery Immunization Enhances HIV Neutralizing Antibody and Germinal Center Responses via Modulation of Immunodominance. Cell 177, 1153–1171.e1128 (2019).

13. Lee, J.H. et al. Long-primed germinal centres with enduring affinity maturation and clonal migration. Nature 609, 998–1004 (2022).

14. Victora, G.D. & Nussenzweig, M.C. Germinal centers. Annu Rev Immunol 30, 429–457 (2012).

15. Tam, H.H. et al. Sustained antigen availability during germinal center initiation enhances antibody responses to vaccination. Proc Natl Acad Sci U S A 113, E6639–E6648 (2016).

16. Shlomchik, M.J. & Weisel, F. Germinal center selection and the development of memory B and plasma cells. Immunol Rev 247, 52–63 (2012).

17. Mesin, L., Ersching, J. & Victora, G.D. Germinal Center B Cell Dynamics. Immunity 45, 471–482 (2016).

18. Gitlin, A.D., Shulman, Z. & Nussenzweig, M.C. Clonal selection in the germinal centre by regulated proliferation and hypermutation. Nature 509, 637–640 (2014).

19. Gitlin, A.D., et al. HUMORAL IMMUNITY. T cell help controls the speed of the cell cycle in germinal center B cells. Science 349, 643–646 (2015).

20. Cyster, J.G. & Allen, C.D.C. B Cell Responses: Cell Interaction Dynamics and Decisions. Cell 177, 524–540 (2019).

21. Avancena, P. et al. The magnitude of germinal center reactions is restricted by a fixed number of preexisting niches. Proc Natl Acad Sci U S A 118 (2021).

22. Roth, G.A. et al. Designing spatial and temporal control of vaccine responses. Nature Reviews Materials (2021).

23. Ou, B.S., Saouaf, O.M., Baillet, J. & Appel, E.A. Sustained delivery approaches to improving adaptive immune responses. Advanced Drug Delivery Reviews 187, 114401 (2022).

24. Hogenesch, H. Mechanism of immunopotentiation and safety of aluminum adjuvants. Front Immunol 3, 406 (2012).

25. HogenEsch, H. Mechanisms of stimulation of the immune response by aluminum adjuvants. Vaccine 20 **Suppl 3**, S34–39 (2002).

26. Shi, Y., HogenEsch, H. & Hem, S.L. Change in the degree of adsorption of proteins by aluminum-containing adjuvants following exposure to interstitial fluid: freshly prepared and aged model vaccines. Vaccine 20, 80–85 (2001).

27. Weissburg, R.P. et al. Characterization of the MN gp120 HIV-1 vaccine: antigen binding to alum. Pharm Res 12, 1439–1446 (1995).

28. Hutchison, S. et al. Antigen depot is not required for alum adjuvanticity. FASEB J 26, 1272–1279 (2012).

29. Noe, S.M., Green, M.A., HogenEsch, H. & Hem, S.L. Mechanism of immunopotentiation by aluminum-containing adjuvants elucidated by the relationship between antigen retention at the inoculation site and the immune response. Vaccine 28, 3588–3594 (2010).

30. Appel, E.A. et al. Self-assembled hydrogels utilizing polymer-nanoparticle interactions. Nat Commun 6, 6295 (2015).

31. Gale, E.C. et al. Hydrogel-Based Slow Release of a Receptor-Binding Domain Subunit Vaccine Elicits Neutralizing Antibody Responses Against SARS-CoV-2. Adv Mater 33, e2104362 (2021).

32. Jons, C.K. et al. Yield-Stress and Creep Control Depot Formation and Persistence of Injectable Hydrogels Following Subcutaneous Administration. Advanced Functional Materials, 2203402 (2022).

33. Meany, E.L. et al. Generation of an inflammatory niche in a hydrogel depot through recruitment of key immune cells improves efficacy of mRNA vaccines. Sci Adv 11, eadr2631 (2025).

34. Ou, B.S. et al. Nanoparticle-Conjugated Toll-Like Receptor 9 Agonists Improve the Potency, Durability, and Breadth of COVID-19 Vaccines. ACS Nano 18, 3214–3233 (2024).

35. Roth, G.A. et al. Injectable Hydrogels for Sustained Codelivery of Subunit Vaccines Enhance Humoral Immunity. ACS Central Science 6, 1800–1812 (2020).

36. Yan, J. et al. A Regimen Compression Strategy for Commercial Vaccines Leveraging an Injectable Hydrogel Depot Technology for Sustained Vaccine Exposure. Advanced Therapeutics 7 (2024).

37. Saouaf, O.M. et al. Sustained Vaccine Exposure Elicits More Rapid, Consistent, and Broad Humoral Immune Responses to Multivalent Influenza Vaccines. Adv Sci (Weinh) 12, e2404498 (2025).

38. Roth, G.A. et al. Prolonged Codelivery of Hemagglutinin and a TLR7/8 Agonist in a Supramolecular Polymer–Nanoparticle Hydrogel Enhances Potency and Breadth of Influenza Vaccination. ACS Biomaterials Science & Engineering (2021).

39. Saouaf, O.M. et al. Modulation of injectable hydrogel properties for slow co-delivery of influenza subunit vaccine components enhance the potency of humoral immunity. J Biomed Mater Res A 109, 2173–2186 (2021).

40. Ou, B.S. et al. Broad and Durable Humoral Responses Following Single Hydrogel Immunization of SARS-CoV-2 Subunit Vaccine. Advanced Healthcare Materials 12 (2023).

41. Böhnert, V., et al. A cGAMP-Containing Hydrogel for Prolonged SARS-CoV-2 Receptor-Binding Domain Subunit Vaccine Exposure Induces a Broad and Potent Humoral Response. Adv Nanobiomed Res 5 (2025).

42. Arunachalam, P.S. et al. T cell-inducing vaccine durably prevents mucosal SHIV infection even with lower neutralizing antibody titers. Nat Med 26, 932–940 (2020).

43. Arunachalam, P.S. et al. A comparative immunological assessment of multiple clinical-stage adjuvants for the R21 malaria vaccine in nonhuman primates. Sci Transl Med 16, eadn6605 (2024).

44. Lee, A. et al. A molecular atlas of innate immunity to adjuvanted and live attenuated vaccines, in mice. Nat Commun 13, 549 (2022).

45. Kasturi, S.P., et al. 3M-052, a synthetic TLR-7/8 agonist, induces durable HIV-1 envelope-specific plasma cells and humoral immunity in nonhuman primates. Sci Immunol 5 (2020).

46. Yu, A.C. et al. Physical networks from entropy-driven non-covalent interactions. Nature Communications 12 (2021).

47. Yu, A.C., Smith, A.A.A. & Appel, E.A. Structural considerations for physical hydrogels based on polymer–nanoparticle interactions. Molecular Systems Design & Engineering 5, 401–407 (2020).

48. Lopez Hernandez, H., Souza, J.W. & Appel, E.A. A Quantitative Description for Designing the Extrudability of Shear-Thinning Physical Hydrogels. Macromolecular Bioscience 21, 2000295 (2021).

49. Eckman, N. & Appel, E.A. Crosslink Dynamics Control Injection Force and Flow Profiles of Non-Covalent Gels. Macromolecules 58, 6350–6358 (2025).

50. Mann, J.L., Yu, A.C., Agmon, G. & Appel, E.A. Supramolecular polymeric biomaterials. Biomater Sci 6, 10–37 (2017).

51. Appel, E.A., Del Barrio, J., Loh, X.J. & Scherman, O.A. Supramolecular polymeric hydrogels. Chem Soc Rev 41, 6195 (2012).

52. Webber, M.J., Appel, E.A., Meijer, E.W. & Langer, R. Supramolecular biomaterials. Nat Mater 15, 13–26 (2016).

53. Correa, S. et al. Injectable Nanoparticle-Based Hydrogels Enable the Safe and Effective Deployment of Immunostimulatory CD40 Agonist Antibodies. Advanced Science, 2103677 (2022).

54. Jons, C.K. et al. Ultrahigh-concentration biologic therapeutics enabled by spray drying with a glassy surfactant excipient. Sci Transl Med 17, eadv6427 (2025).

55. Hill, R.L. et al. Comparison of drug delivery with autoinjector versus manual prefilled syringe and between three different autoinjector devices administered in pig thigh. Med Devices (Auckl) 9, 257–266 (2016).

56. Moon, J.J. et al. Enhancing humoral responses to a malaria antigen with nanoparticle vaccines that expand Tfh cells and promote germinal center induction. Proc Natl Acad Sci U S A 109, 1080–1085 (2012).

57. Moyer, T.J. et al. Engineered immunogen binding to alum adjuvant enhances humoral immunity. Nat Med 26, 430–440 (2020).

58. Rodrigues, K.A. et al. Phosphate-mediated coanchoring of RBD immunogens and molecular adjuvants to alum potentiates humoral immunity against SARS-CoV-2. Sci Adv 7, eabj6538 (2021).

59. Ou, B.S. et al. Saponin nanoparticle adjuvants incorporating Toll-like receptor agonists drive distinct immune signatures and potent vaccine responses. Sci Adv 10, eadn7187 (2024).

60. Hu, M. et al. Clonal composition and persistence of antigen-specific circulating T follicular helper cells. Eur J Immunol 53, e2250190 (2023).

61. Arunachalam, P.S. et al. Durability of immune responses to mRNA booster vaccination against COVID-19. J Clin Invest 133 (2023).

62. Hu, M. et al. Altered baseline immunological state and impaired immune response to SARS-CoV-2 mRNA vaccination in lung transplant recipients. Cell Rep Med 6, 102050 (2025).

63. Arunachalam, P.S. et al. Durable protection against the SARS-CoV-2 Omicron variant is induced by an adjuvanted subunit vaccine. Sci Transl Med 14, eabq4130 (2022).

64. Feng, Y. et al. Broadly neutralizing antibodies against sarbecoviruses generated by immunization of macaques with an AS03-adjuvanted COVID-19 vaccine. Sci Transl Med 15, eadg7404 (2023).

65. Grigoryan, L., et al. AS03 adjuvant enhances the magnitude, persistence, and clonal breadth of memory B cell responses to a plant-based COVID-19 vaccine in humans. Sci Immunol 9, eadi8039 (2024).

66. Arunachalam, P.S. et al. Durable protection against the SARS-CoV-2 Omicron variant is induced by an adjuvanted subunit vaccine. Science Translational Medicine 14, eabq4130 (2022).

67. Andrews, N. et al. Covid-19 Vaccine Effectiveness against the Omicron (B.1.1.529) Variant. N Engl J Med 386, 1532–1546 (2022).

68. Hsieh, C.L. et al. Structure-based design of prefusion-stabilized SARS-CoV-2 spikes. Science 369, 1501–1505 (2020).

69. Gibson, D.G. et al. Enzymatic assembly of DNA molecules up to several hundred kilobases. Nat Methods 6, 343–345 (2009).

70. Xie, X. et al. An Infectious cDNA Clone of SARS-CoV-2. Cell Host Microbe 27, 841–848 e843 (2020).

71. Pegu, A., et al. Durability of mRNA-1273 vaccine-induced antibodies against SARS-CoV-2 variants. Science 373, 1372–1377 (2021).

72. Edara, V.V. et al. Infection and Vaccine-Induced Neutralizing-Antibody Responses to the SARS-CoV-2 B.1.617 Variants. N Engl J Med 385, 664–666 (2021).

73. Edara, V.V., et al. mRNA-1273 and BNT162b2 mRNA vaccines have reduced neutralizing activity against the SARS-CoV-2 omicron variant. Cell Rep Med 3, 100529 (2022).

74. Vanderheiden, A. et al. Development of a Rapid Focus Reduction Neutralization Test Assay for Measuring SARS-CoV-2 Neutralizing Antibodies. Curr Protoc Immunol 131, e116 (2020).

75. Katzelnick, L.C. et al. Viridot: An automated virus plaque (immunofocus) counter for the measurement of serological neutralizing responses with application to dengue virus. PLoS Negl Trop Dis 12, e0006862 (2018).

