## Supporting Information for "Sustained Delivery of a SARS-CoV-2 Subunit Vaccine in An Adjuvanted Hydrogel Depot Enhances Vaccine Responses in Nonhuman Primates"

Ben S. Ou *et al.*

**This PDF file includes:**

Figures S1 to S11
Tables S1

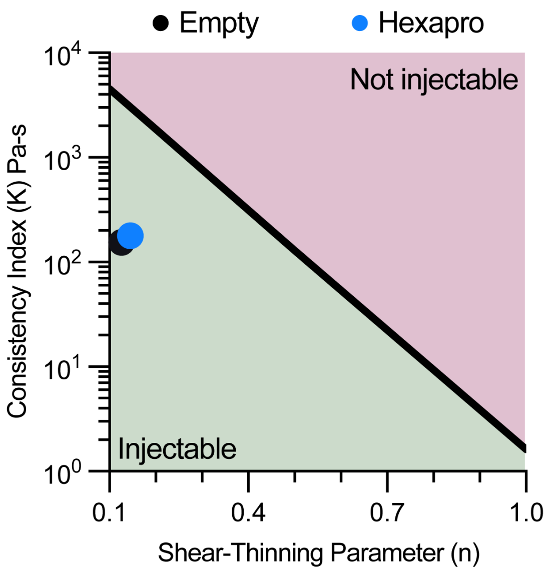


**Figure S1. Clinically Relevant Extrudability.** Ashby-style plot demonstrating the injectability of PNP hydrogels, with or without cargo, under clinically relevant constraints. Based on methodology from Lopez Hernandez et al.[1]

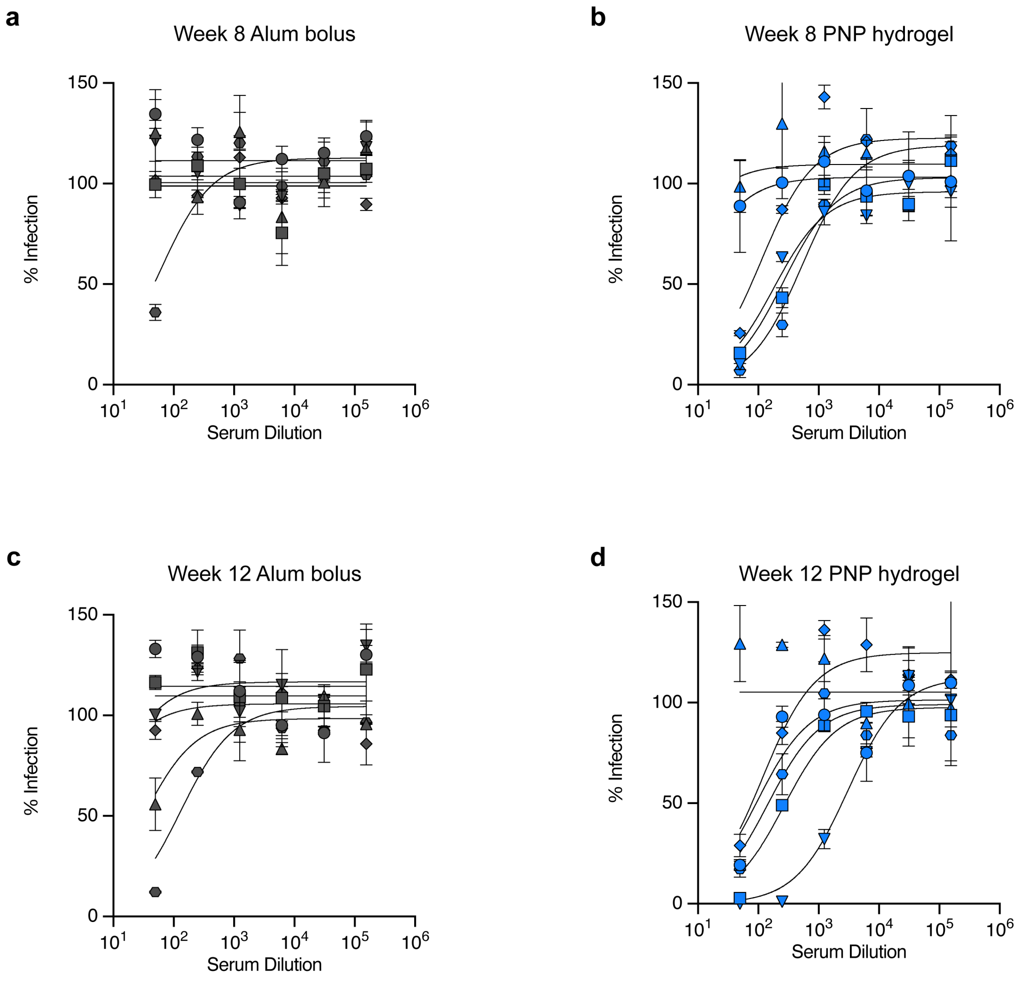


**Figure S2. Neutralizing Antibodies in Mice Post-Hexapro Vaccines. (a-d)** Percent infectivity for all vaccine treatments at a range of Week 8 and Week 12 serum dilutions as determined by a SARS-CoV-2 spike-pseudotyped viral neutralization assay.

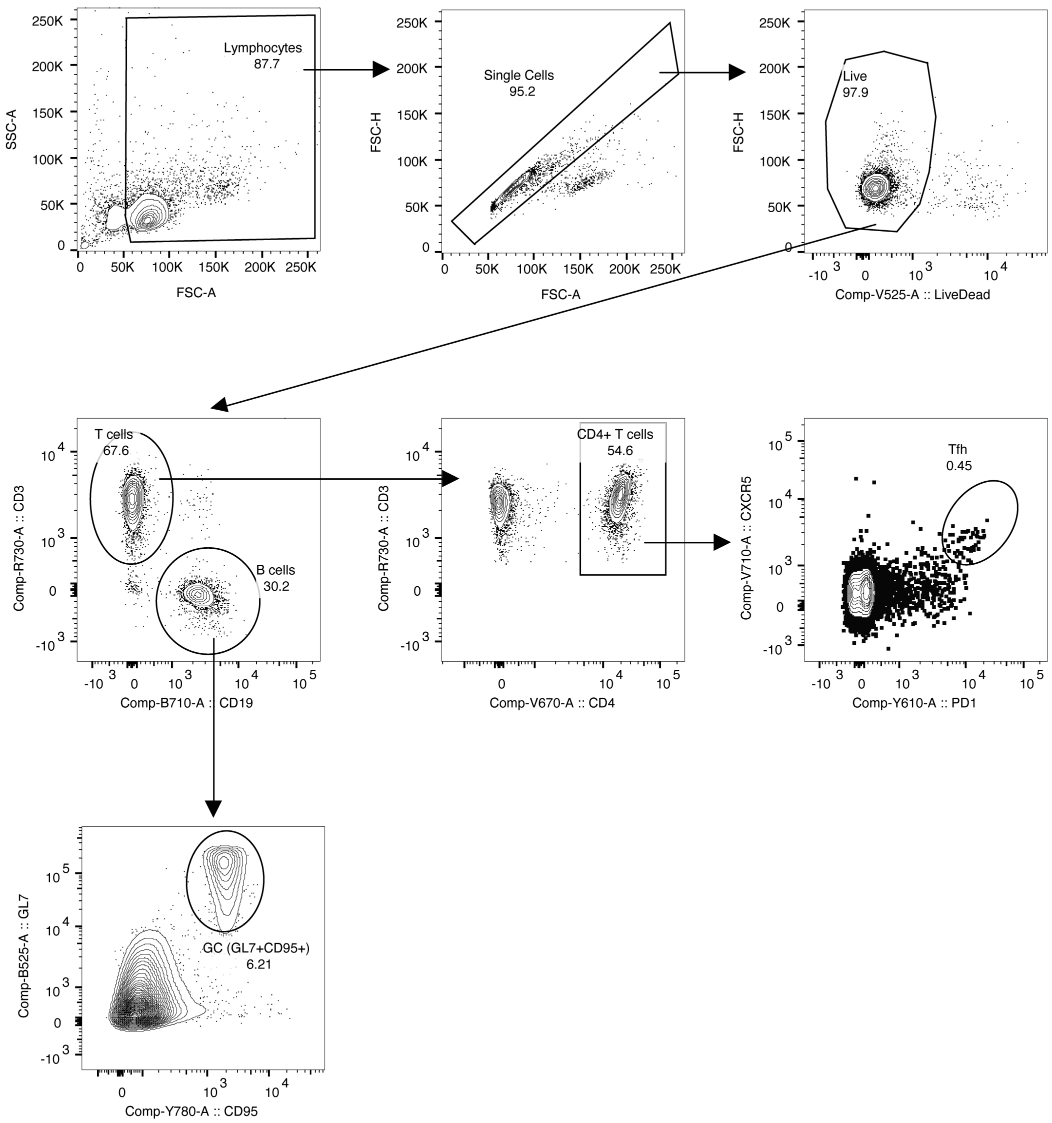


**Figure S3. Lymph Node Flow Cytometry Gating in Mice.** Gating strategy for lymph nodes to assess germinal center reactions. A representative sample of the PNP hydrogel treatment group was used.

**a**

| Score | Grade | Edema | Erythema |
|---|---|---|---|
| 0 | None | No swelling | Normal pink |
| 1 | Minimal | Slight swelling, indistinct border | Light pink, indistinct |
| 2 | Mild | Defined swelling, distinct border | Bright pink, distinct |
| 3 | Moderate | Defined swelling, raised border (<1mm) | Light red, distinct |
| 4 | Severe | Pronounced swelling, raised border (> or = 1mm) | Dark red, pronounced |

**b**

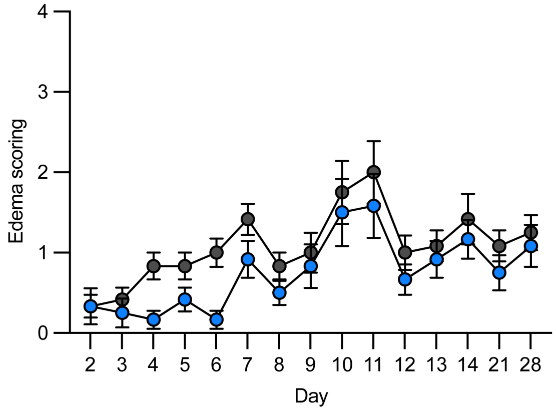


**c**

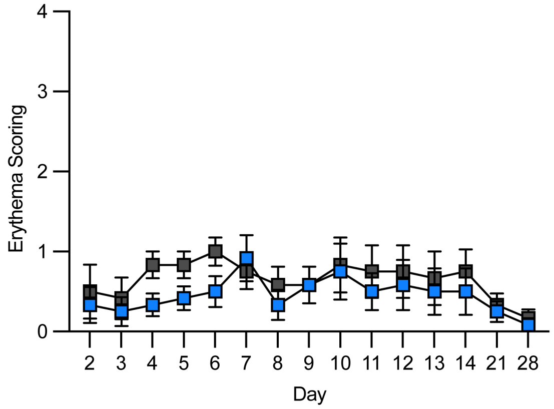


**Figure S4. Injection-Site Reactions Following Hexapro Vaccines. (a)** Clinical edema and erythema scores were assessed daily for the first two weeks, then weekly until Week 4 following RM immunizations with either Alum bolus or PNP hydrogel Hexapro vaccines. **(b)** Clinical edema scores over 28 days period. **(c)** Clinical erythema scores over 28 days period. Data are shown as mean ± SEM (n = 6). The *p* values listed were determined by the unpaired Mann-Whitney test.

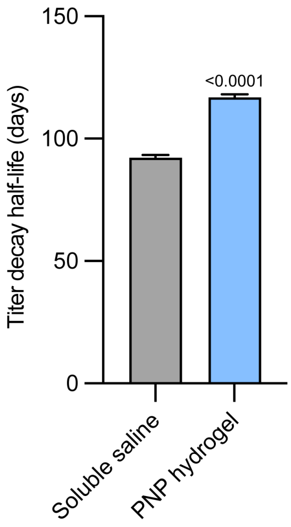


**Figure S5. Antibody decay half-life post-boost.** Decay half-life values derived from parametric bootstrapping of titers (n=1000 simulations) following each treatment group's post-boost peak. Data are shown as bars with mean ± SEM (n = 6). The *p* values listed were determined by the (a) unpaired Mann-Whitney test or (b) Unpaired t test with Welch's correction.

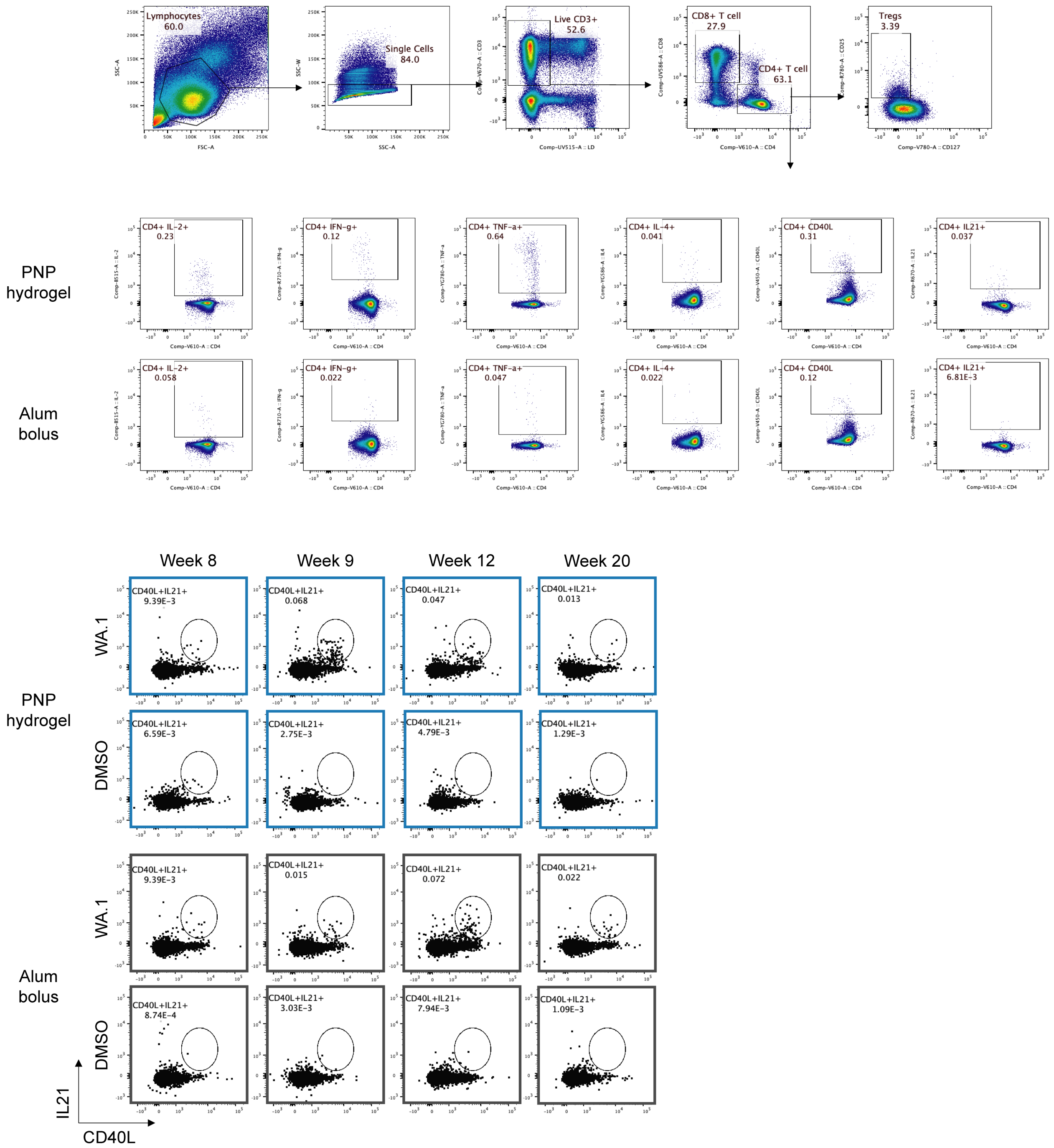


**Figure S6. T Cell Intercellular Staining Flow Cytometry Gating in NHP.** Gating strategy for PBMCs to assess T cell population vis intercellular staining.

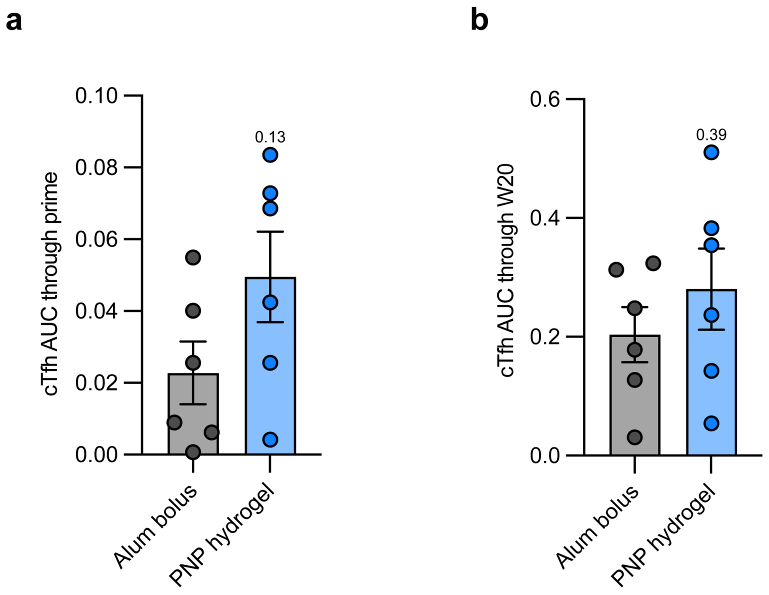


**Figure S7. Area under the curve of circulating T follicular helper cells (cTfhs).** Values determined **(a)** during priming or **(b)** over 20 weeks. Data are shown as bars with mean ± SEM (n = 6). The *p* values listed were determined by the unpaired Mann-Whitney test.

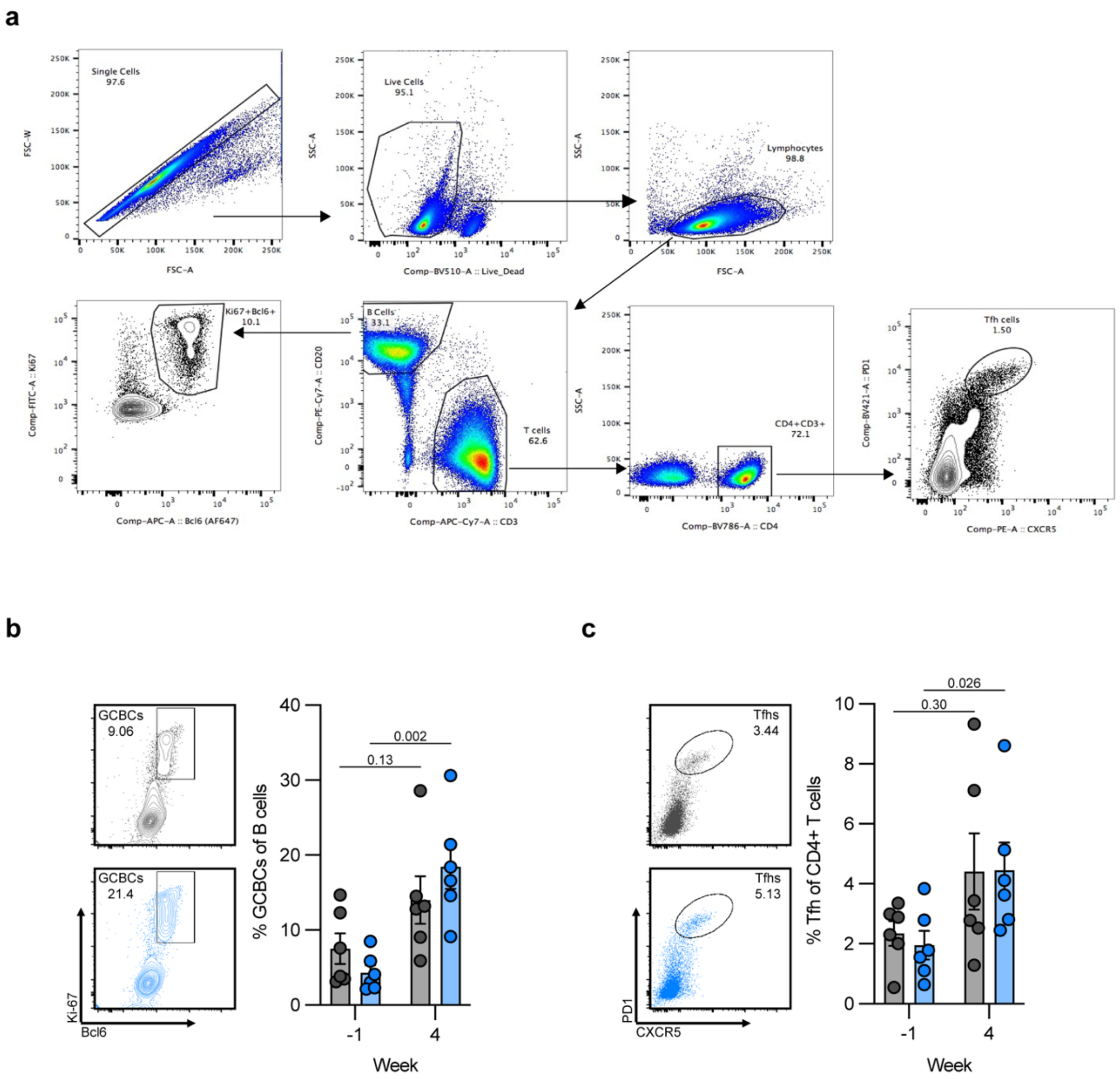


**Figure S8. Lymph Node Flow Cytometry in NHP. (a)** Gating strategy for lymph nodes to assess germinal center reactions. **(b)** Representative flow plots and percentages of germinal center B cell (GCBC) of each vaccine group pre- and 4 weeks after inoculation. **(c)** Representative flow plots and percentage of germinal center Tfh cell (GC-Tfh) of each vaccine group pre- and 4 weeks after inoculation.

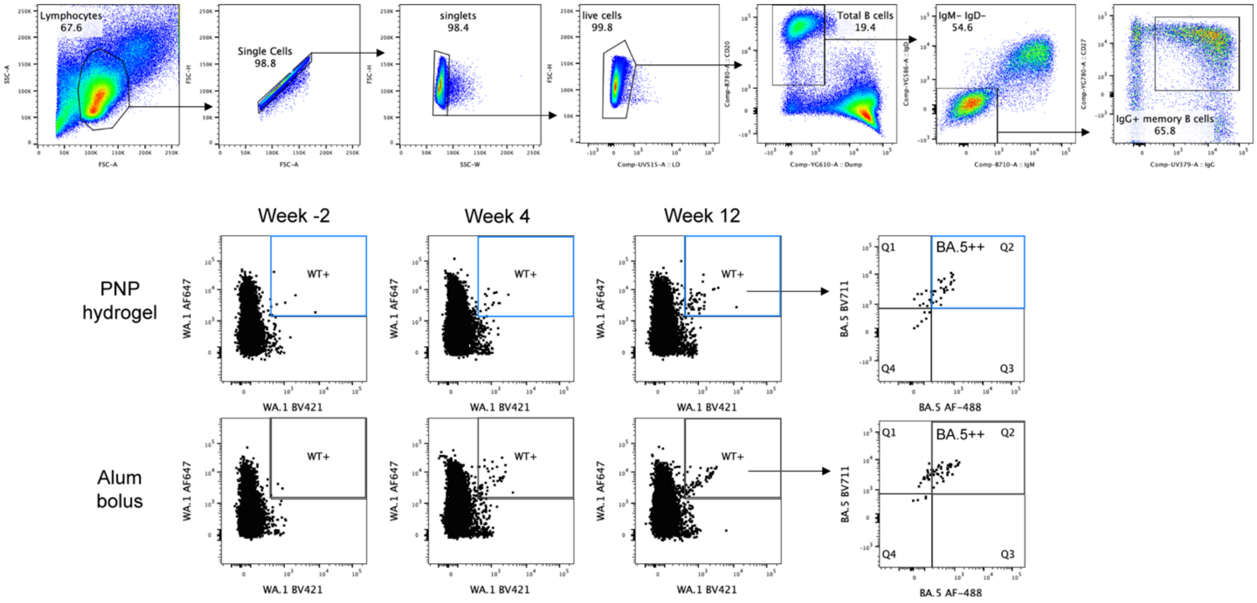


**Figure S9. Memory B cell Flow Cytometry Gating in NHP.** Gating strategy for PBMCs to assess Memory B cell population.

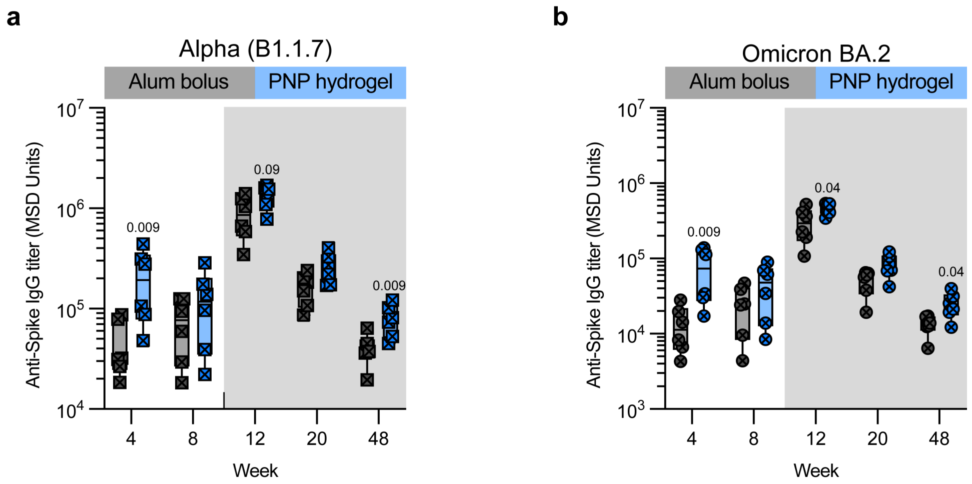


**Figure S10. Binding antibody titers**. Titers against **(a)** Alpha (B1.1.7) and **(b)** Omicron BA.2 variants following each vaccine treatment. Data are shown as box-and-whiskers plots showing median, interquartile (box), and range (whiskers) (n = 6). The *p* values listed were determined by the unpaired Mann-Whitney test.

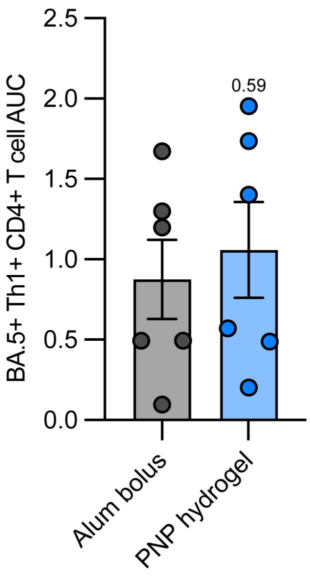


**Figure S11. Area under the curve of Omicron BA.5-specific CD4+ T cell production over 20 weeks.** Data are shown as bars with mean ± SEM (n = 6). The *p* values listed were determined by the unpaired Mann-Whitney test.

**Table S1. Antibody Markers Used for T cell Intercellular Staining Flow Cytometry.**

| | **Marker** | **Vendor** | **Catalog#** | **Clone** |
|---|---|---|---|---|
| **FITC** | IL-2 | Biolegend | 500304 | MQ1-17H12 |
| **PE** | IL-4 | BioLegend | 500810 | MP4-25D2 |
| **PE-CF594** | CD45RA | BD Biosciences | 565419 | 5H9 |
| **PE-Cy7** | TNFa | E-Bioscience | 25-7349-82 | Mab11 |
| **BV421** | CD40L | Biolegend | 310824 | 24-31 |
| **BV506** | TCRgd | Biolegend | 331220 | B1.1 |
| **BV 605** | CD4 | Biolegend | 317438 | OKT4 |
| **BV650** | CD3 | BD Biosciences | 563916 | SP34-2 |
| **BV711** | CCR7 | Biolegend | 353228 | G043H7 |
| BV785 | CD127 | Biolegend | 351330 | A019D5 |
| **APC** | IL-21 | BioLegend | 513008 | 3A3-N2 |
| **A700** | IFNg | Biolegend | 502520 | 4S.B3 |
| **APC-Cy7** | CD25 | Biolegend | 302614 | BC96 |
| **BUV496** | LD | Biolegend | 423108 | |
| **BUV563** | CD8 | BD Biosciences | 612914 | RPA-T8 |
| **BUV737** | CCR6 | BD Biosciences | 612780 | 11A9 |
| **BUV805** | CD69 | BD Biosciences | 748763 | FN50 |

**References**

1. Lopez Hernandez, H., Souza, J.W. & Appel, E.A. A Quantitative Description for Designing the Extrudability of Shear-Thinning Physical Hydrogels. *Macromolecular Bioscience* **21**, 2000295 (2021).
